# 3D Additive Manufactured Composite Scaffolds with Antibiotic-loaded Lamellar Fillers for Bone Infection Prevention and Tissue Regeneration

**DOI:** 10.1101/2020.05.07.082495

**Authors:** María Cámara-Torres, Stacy Duarte, Ravi Sinha, Ainhoa Egizabal, Noelia Álvarez, Maria Bastianini, Michele Sisani, Paolo Scopece, Marco Scatto, Alessandro Bonetto, Antonio Marcomini, Alberto Sanchez, Alessandro Patelli, Carlos Mota, Lorenzo Moroni

## Abstract

Bone infections following open bone fracture or implant surgery remain a challenge in the orthopedics field. In order to avoid high doses of systemic drug administration, optimized local antibiotic release from scaffolds is required. 3D additive manufactured (AM) scaffolds made with biodegradable polymers are ideal to support bone healing in non-union scenarios and can be given antimicrobial properties by the incorporation of antibiotics. In this study, ciprofloxacin and gentamicin intercalated in the interlamellar spaces of magnesium aluminum layered double hydroxides (MgAl) and α-zirconium phosphates (ZrP), respectively, are dispersed within a thermoplastic polymer by melt compounding and subsequently processed via high temperature melt extrusion AM (∼190 °C) into 3D scaffolds. The inorganic fillers enable a sustained antibiotics release through the polymer matrix, controlled by antibiotics counterions exchange or pH conditions. Importantly, both antibiotics retain their functionality after the manufacturing process at high temperatures, as verified by their activity against both Gram + and Gram − bacterial strains. Moreover, scaffolds loaded with filler-antibiotic do not impair human mesenchymal stromal cells osteogenic differentiation, allowing matrix mineralization and the expression of relevant osteogenic markers. Overall, these results suggest the possibility of fabricating dual functionality 3D scaffolds via high temperature melt extrusion for bone regeneration and infection prevention.

## 1. Introduction

In the orthopedics field, surgical procedures involving fracture stabilizations or implants can develop infections in up to 5% of the cases, while this rate rises up to 50% in open bone fracture scenarios. ^[1]^ Bacterial infections can trigger osteomyelitis, an inflammatory reaction in bone that can ultimately lead to bone destruction or osteolysis. ^[2]^ Moreover, bacteria can invade and survive within osteoblasts slowing down or preventing new bone formation, or create biofilms, which present a challenge to the antibiotic therapy, due to reduced antimicrobial susceptibility leading to prolonged infection. ^[3]^ The standard medical strategy to overcome osteomyelitis involves the debridement of infected bone and soft tissue and the removal of the implant or fixation device, accompanied by the systemic administration of antibiotics from 6 to 16 weeks depending on the severity of the infection. ^[4]^ Typically, high antibiotic concentrations are administered systemically to achieve effective concentrations at the infected site, which can lead to the development of resistant bacterial strains, as well as cell toxicity and adverse effects in the bone regeneration process. ^[5]^ Since the removal of necrotic bone tissue commonly leaves a critical size defects, antibiotic-laden poly(methyl methacrylate) (PMMA) cements or beads were designed to act both as bone spacer and as local antibiotic delivery systems, maximizing target tissue concentrations and minimizing toxicity risks. ^[6]^ Despite being one of the gold standard approaches, PMMA cements possess a series of disadvantages including lack of biodegradability, requirement of a second surgery to replace it by other material that allows bone regeneration. Furthermore PMMA cements present a suboptimal drug elution kinetics generally associated to a non-reproducible antibiotic-polymer mixing procedure, and due to the exothermic polymerization of PMMA, only a restricted amount of heat-stable antibiotics can be incorporated. ^[7]^ With the recent development in the field of biomaterials and bone tissue engineering, a number of alternative carriers have been designed to allow biodegradability and the capability of delivering locally a wider variety of antimicrobial agents. These carriers are mainly collagen, chitosan, calcium sulfate, calcium phosphates, demineralized bone, as well as synthetic polymers, such as polyurethanes, polyanhydrides, poly(lactic acid) (PLA) and poly(lactides-*co*-glycolides) (PLGA). ^[8]^ However, processing these materials into 3D scaffolds using conventional fabrication techniques, such as gas foaming/particulate leaching, freeze-drying or phase separation, can result in constructs that lack reproducibility or the structural integrity and mechanical properties needed to stabilize a non-union defect, making them also not fully functional candidates for facilitating bone healing following infection management.

In the past decade, additive manufacturing (AM) and, more specifically, melt extrusion-based techniques, have emerged as one of the most appealing technologies to produce scaffolds for bone tissue engineering. ^[9]^ This method allows for the reproducible fabrication of patient-personalized 3D scaffolds, from a variety of biocompatible and biodegradable thermoplastic polymers, with an interconnected pore network, high porosity, and optimum mechanical properties for bone regeneration. Osteoinductive and antibacterial scaffolds have recently been developed by combining AM and drug delivery. Antimicrobial peptides, ^[10]^ or other antimicrobial compounds, such as silver, ^[11]^ bioactive glass, ^[12]^ or quaternized chitosan, ^[13]^ have been incorporated onto the surface of 3D AM scaffolds. These alternative compounds have been used to avoid the use of antibiotics due to their potential cell toxicity and bacterial resistance risks. Yet, the *in vivo* efficacy of these substitutes compared to conventional antibiotic therapies is still unproven. Therefore, as a common strategy, antibiotics have been adsorbed on the surface of AM scaffolds. However, surface adsorption via the immersion of the scaffold in an antibiotic solution has shown a limit retention of the loaded antibiotic, amount that correlates to the scaffold surface area, resulting in poor loading efficiency and leading to the burst release of the antibiotic. ^[14]^ In order to increase the loading efficiency and allow for a more sustained drug release, different strategies have been explored, such as porogen leaching to create microporosity in the scaffold’s filaments, hydrogel coatings, or loading of antibiotics within degradable polymeric microparticles. ^[14–15]^ The combination of several of these loading methods in the same scaffold has also been considered to allow the incorporation of different antibiotics and to obtain a sequential release profile for a more potent therapy. ^[16]^ However, in spite of enhancing the scaffold antimicrobial properties, these multi-step post-modification approaches are complex and laborious. Thus, the direct incorporation of the antibiotic within the scaffold material would be desired to simplify the fabrication process, as well as to improve the loading capability and release profile. With the exception of melt extrusion at low temperatures, ^[17]^ the incorporation of antibiotics in the scaffold material is not frequently investigated due to the sensitivity of bioactive molecules to temperature. This in turn limits the polymer choice, as the majority of thermoplastic polymers for biomedical applications require long exposure to high processing temperatures (∼175 – 220 °C), ^[18]^ which are above the thermal stability limits of most commonly used antibiotics (∼ 120 °C, depending on exposure time and antibiotic type). ^[19]^ Moreover, most of current delivery systems lack the optimum release kinetics of the cargo. A desirable sustained drug release should ensure that an adequate antibiotic concentrations is delivered on site, above the minimal inhibitory concentration (MIC) values and over a sufficient time to cover the critical window period post-surgery, followed by a sustained release at an effective level for inhibiting the occurrence of a latent infection. ^[20]^

Here, we introduce a novel approach that allows the direct fabrication of drug loaded scaffolds via high temperature melt extrusion AM. The composite scaffolds consist of the biocompatible and biodegradable copolymer poly(ethyleneoxideterephthalate)/poly(butyleneterephthalate) (PEOT/PBT) and the inorganic layered compounds magnesium aluminium layered double hydroxide (MgAl) or α-Zirconium phosphate (ZrP), which contain antibiotics intercalated within their lamellar structures. Ciprofloxacin (CFX) and gentamicin (GTM), two commonly used antibiotics to treat orthopedic infections, are used as model anionic and cationic molecules to be intercalated within the positively charged MgAl and negatively charged ZrP fillers, respectively. We evaluate the dual functionality of these scaffolds as i) local antibiotic delivery systems against *Staphylococcus epidermidis* (Gram +) and *Pseudomonas aeruginosa (Gram* −), identified as two of the major responsible for implant associated infections ^[21]^, and ii) their potential to support bone tissue formation.

## 2. Results and discussion

### 2.1 Scaffolds fabrication and characterization

Although the sustained release of drugs from MgAl ^[22]^ and ZrP ^[23]^ has been previously reported, the combination of antibiotic-loaded fillers with polymers has not been carefully examined in terms of release kinetics, antimicrobial functionality or ensurance of no negative effect on the bone regeneration response. Most importantly, no previous studies have reported their processing into 3D AM scaffolds, but only as implant coatings ^[24]^ or incorporated on electrospun scaffolds. ^[25]^ Here, 3D polymer composite scaffolds with filler-antibiotic concentrations up to 20 wt% (MgAl-CFX_s or ZrP-GTM_s) were successfully fabricated for the first time via melt extrusion AM of previously prepared melt-blended composite pellets (MgAl-CFX_p or ZrP-GTM_p) (**Figure 1** and **Figure S1**). All scaffolds exhibited interconnected macropores, and experimental filament diameters, and X-Y and Z pores sizes consistently matched the theoretical values (**Figure S2**).

**Figure 1.**
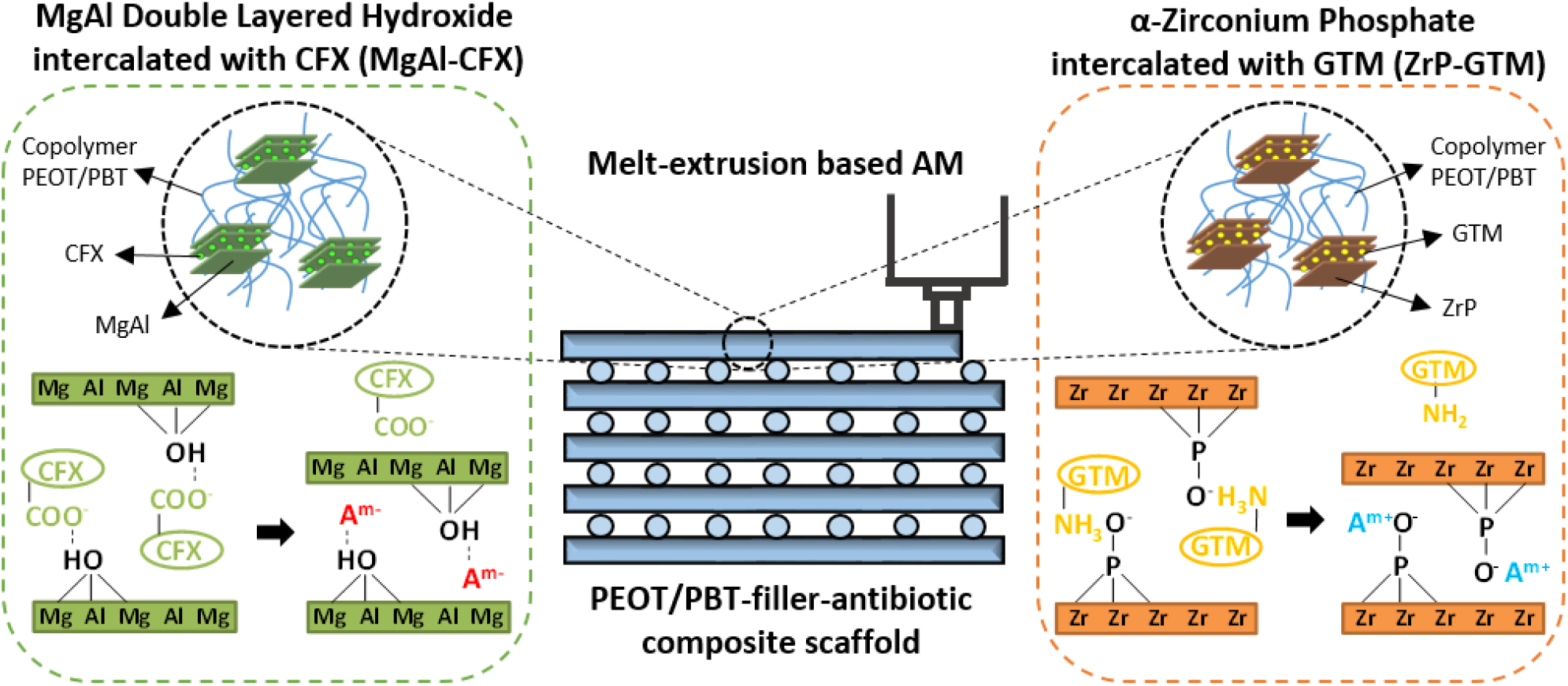
Schematic representation of the antibacterial composite scaffolds (PEOT/PBT-filler-antibiotic) fabricated by melt-extrusion AM. These polymeric composite scaffolds contain either MgAl intercalated with CFX (MgAl-CFX), or ZrP intercalated with GTM (ZrP-GTM). When submerged in an eluent solution, anions can exchange with CFX, which diffuses out of the filler and polymer matrix into the eluent. On the other hand, cations can exchange with GTM in the ZrP-GTM based system, allowing the diffusion of GTM out of the filler and polymer matrix. The release of antibiotics confer antimicrobial activity to the scaffolds.

As it is observed in **Figure 2**, backscattered electron (BSE) images confirm an increasing amount of filler (visible as brighter points) with increasing filler-antibiotic concentrations on the surface and cross section of the scaffolds filaments. Using energy dispersive X-ray spectroscopy (EDS) the fillers were discriminated as MgAl and ZrP containing particles and the distinctive atomic elements of the CFX (F and N) and GTM (S and N) molecules were found to co-localize with their respective lamellar particles (**Figure 2C-D**). This strongly indicates that the filler and antibiotic were not separated in the filaments, but showed a preserved filler structure after the melt-blending and AM process. Moreover, particle size was maintained for both types of fillers within the after-synthesis values during the processing steps, with a slight size reduction during the AM process, likely due to the higher shear forces in the print-head screw (**Figure S3**).

**Figure 2.**
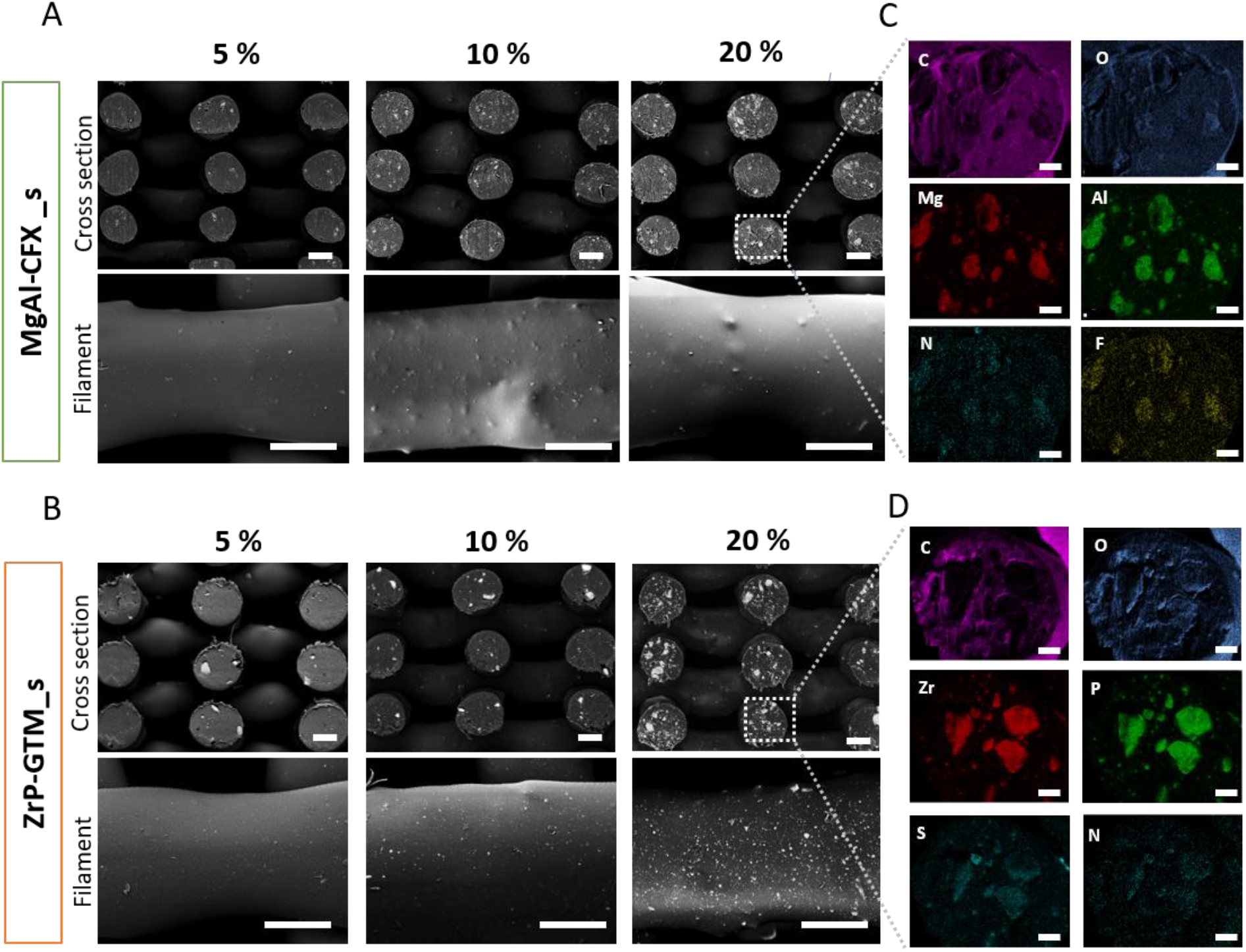
Filler distribution and antibiotics localization on composite scaffolds. BSE micrographs of scaffolds cross sections and filaments surface depicting filler distribution (as white points) on (A) MgAl-CFX_s and (B) ZrP-GTM_s at different filler-antibiotic concentrations. Scale bars 250 μm. EDAX elemental mapping of representative fiber cross section of (C) 20 % MgAl-CFX_s and (D) 20 % ZrP-GTM_s. Scale bars 50 μm.

### 2.2 *In vitro* antibiotic release

In order to control the kinetics of antibiotic release from the MgAl-CFX and ZrP-GTM composite systems, we explored several approaches including varying the i) filler-antibiotic concentration, ii) the eluent pH and iii) the eluent ionic concentration. As demonstrated in **Figure 3A**, by increasing the MgAl-CFX concentration within MgAl-CFX_s, larger amounts of CFX were released over time. A similar trend was also observed in the release profile of GTM from ZrP-GTM_s (**Figure 3B**). This tendency demonstrates the possibility of designing a customized scaffold, which releases the desired amount of antibiotic over time, by tuning the filler-antibiotic concentration mixed with the polymer matrix. Although both systems demonstrated a filler-antibiotic concentration-dependent release profile, significantly different release kinetics were observed on MgAl-CFX_s and ZrP-GTM_s. While the release of CFX from MgAl-CFX_s was sustained over the course of the evaluated time (1 month), there was a GTM burst release from ZrP-GTM_s during the first 24h followed by a slower release up to the completion of the study (**Figure 3 A-B**). When active molecules are fixed into the inorganic lamellae of layered double hydroxides (LDH), such as MgAl, their de-intercalation mechanism occurs mainly by anion exchange, a process in which the intercalated active molecule and free anions in the elution solution can be exchanged via a concentration gradient. ^[26]^ On the other hand, the release of active molecules from metal hydrogen phosphates, such as ZrP, can occur not only by ion exchange (in this case, of cations), but also via the disruption of acid-base interactions between the active molecule and the filler, which rely on a pH switch in the environment about the acid dissociation constant (pKa) of the intercalated molecule ^[26]^ (**Figure 1**). A series of factors are involved in the release kinetics of antibiotics when the lamellar particles are dispersed within a polymeric matrix: i) the permeation and diffusion of water molecules carrying counter ions through the polymer matrix; ii) the intra-particle diffusion of water, ions and active molecules, which depends on guest-host interactions and determines the rate of de-intercalation, and iii) the diffusion rate of active molecules (antibiotics in this study) from the polymer matrix. ^[27]^ In order to determine the release rate-limiting step, the release profiles of the 20 wt% filler-antibiotic scaffolds, taken as a representative cases, were fitted to various release kinetic models. 7 relevant models were used and the corresponding correlation coefficients are reported in **Figure S4**. Bhaskar model was found to best describe the release profile of CFX from 20% MgAl-CFX_s. This model has previously been used to explain the release of active molecules from LDH systems and indicates that the release of CFX is an intra-particle diffusion controlled process, and therefore, ion exchange rate is the limiting step. ^[28]^ Accordingly, the decreased release rate of CFX over time (**Figure 3A**) is likely due to the formation of a phase boundary between internal zones of the lamellar filler containing intercalated CFX and the external zones, in which anions from dPBS have already replaced the antibiotic, which progressively declines the antibiotic release process. ^[29]^

**Figure 3.**
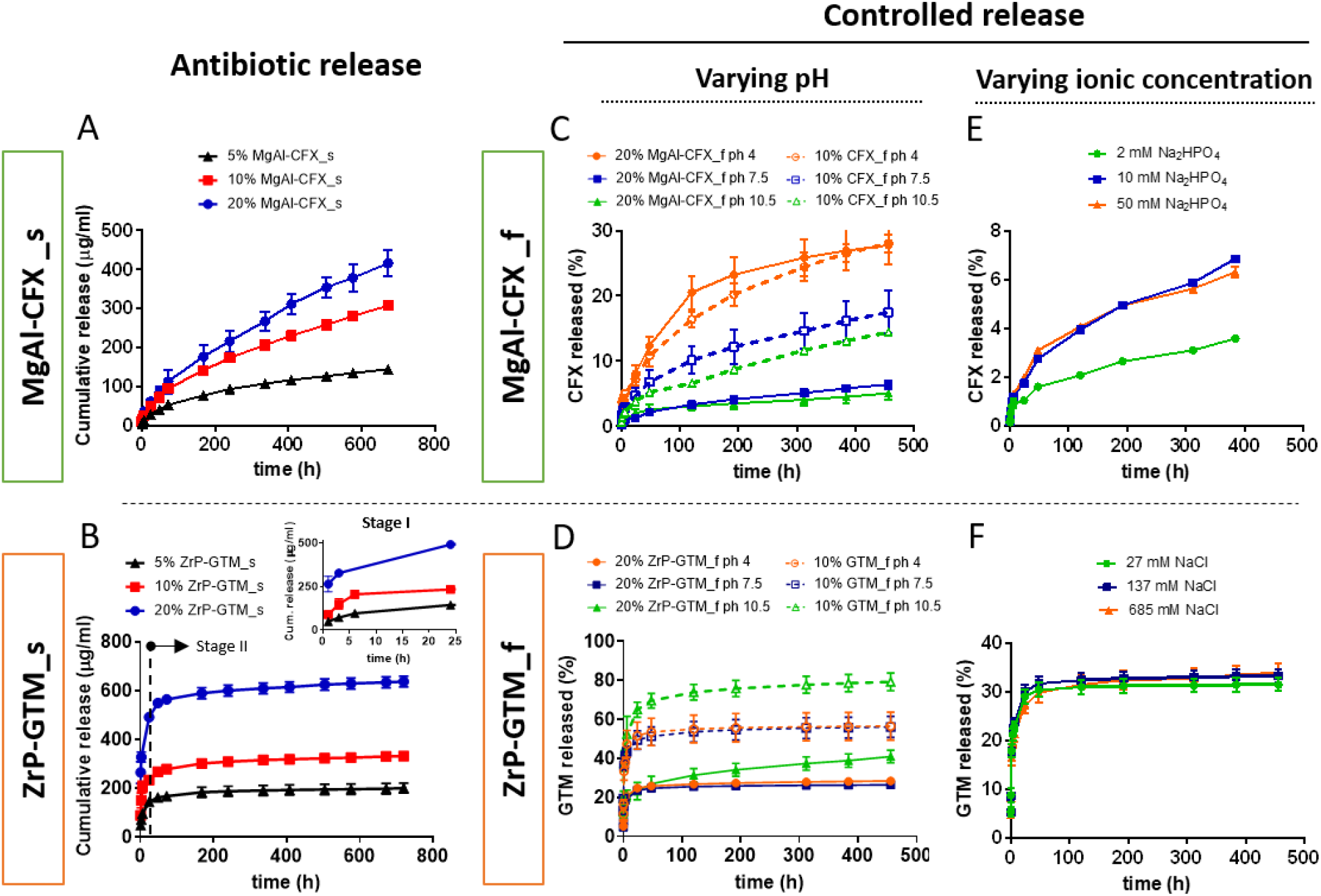
Antibiotic release profiles from MgAl-CFX and ZrP-GTM copolymer composites under different eluent conditions. Cumulative release of (A) CFX from MgAl-CFX_s and (B) GTM from ZrP-GTM_s immersed in dPBS during 4 weeks at 37 °C, proving the filler-antibiotic concentration dependent release. Influence of filler presence and pH of the eluent solution on the cumulative release percentage of (C) CFX and (D) GTM from filler-antibiotic composite films (20% MgAl-CFX_f and 20% ZrP-GTM_f, respectively) and films composed of comparable amounts of antibiotic directly dispersed in the copolymer matrix (10% CFX_f and 10% GTM_f), immersed in buffers at acidic, neutral and basic pH during 3 weeks at 37 °C. Influence of the ionic concentration of the eluent solution on the cumulative release percentage of (E) CFX and (F) GTM from 20% MgAl-CFX_f and 20% ZrP-GTM_f, respectively, during 3 weeks a 37 °C.

To gain more insight into the release mechanism of 20% ZrP-GTM_s, the total progress was separated into two stages: a rapid release stage I (<24h) followed by a slow release stage II (>24h). The modified Freundlich kinetic model, which describes heterogeneous diffusion from flat surfaces of clays via ion exchange, was found to explain the release in Stage I (**Figure S4**). This implies that GTM diffuses into the medium solution via anion exchange during the first 24h, at a fast speed due to the antibiotic high solubility in water-based neutral pH eluents. ^[30]^ Since after the studied period the GTM released did not reach 100% (**Figure S4 A**), it is hypothesized that GTM released in stage I came from the ZrP particles located on the surface of scaffold filaments, after which the release of GTM from internal ZrP particles was hindered by the polymer network, resulting in the plateau-like profile observed in Stage II (**Figure 3B)**. This type of behavior was already reported and is likely due to the low diffusion rate of water molecules through the polymer network, given by the poor water uptake/swellability capability of PEOT/PBT (4% water uptake) at the specific PEO molecular weight (300 kDa) and PEOT:PBT segments ratio (55:45). ^[27, 31]^ In this case, the GTM still incorporated into the system could be released upon polymer degradation, a process that could be potentially speeded up by increasing the molecular weight of the PEO segments of the copolymer or by using a different and faster degrading material. Following the same reasoning, the fact that a plateau was not reached in the CFX release curve from MgAl-CFX_s, and that only ∼ 15-20% of the total CFX was released (**Figure S4 A**) in the analyzed time, suggests that the CFX released during the evaluated period was also originated from the MgAl particles localized on the surface of the filaments. It is plausible that due to the low solubility of CFX at neutral pH, and possibly the stronger filler-antibiotic interactions, the fast release seen in Stage I in 20% ZrP-GTM_s did not happen in 20% MgAl-CFX_s.

#### Effect of filler and pH

In order to investigate the effect of the filler, antibiotic release from filler-antibiotic composite films (20 % MgAl-CFX_f) and films made of the same amount of CFX directly dispersed in the copolymer matrix (10% CFX_f) was compared (**Figure 3C)**. The presence of the filler enabled the maintenance of a lower antibiotic release rate over time. The effect of filler on the sustained release of CFX was further confirmed upon incubation of 20% MgAl-CFX_f in acidic pH (pH = 4) (**Figure 3C**). Here, the acidic attack dissolved the layered MgAl, allowing for a faster release of CFX, whose release was not governed by ion exchange anymore and became comparable to the release of CFX from 10% CFX_f at pH = 4. ^[28c, 32]^ Interestingly, at pH ≥ 7, 10% CFX_f release profile remained unchanged and lower release rates than at pH = 4 were observed, due to lower CFX solubility at this pH range. Similarly, 20% MgAl-CFX_f release profile did not vary at pH ≥ 7, as MgAl remained stable and intact (**Figure 3C**). The role of ZrP on maintaining a more sustained release, especially in the stage I, was also demonstrated for 20% ZrP-GTM_f, when comparing to 10% GTM_f (**Figure 3D**). However, while MgAl in MgAl-CFX_f was affected by acidic pH, the release of GTM from 20% ZrP-GTM_f was affected by alkaline conditions (pH = 10.5), due to the different nature of the system (**Figure 3D**). Considering the isoelectric point of GTM (pKa = 8), ^[33]^ the hydroxyl groups of GTM were deprotonated at pH = 10.5, disrupting the hydrogen bonds with the co-intercalated water molecules as well as producing a negatively charged GTM, which was repelled from the ZrP filler due to the disrupted acid-base electrostatic interactions between the GTM amine groups and the phosphate groups of the ZrP layers. ^[23a]^ It is hypothesized that oxidative degradation of PEOT/PBT, happening at alkaline conditions, ^[31, 34]^ allowed for further increase in the diffusion of GTM from the polymer matrix, which is visible as a slight increase in the release rate of GTM in Stage II on both 20% ZrP-GTM_f and 10% GTM_f release curves at pH =10 (**Figure 3D**).

#### Effect of ionic concentration

The aforementioned results indicate that the GTM release from ZrP-GTM_f in stage II can be controlled by varying the pH (acid base-interactions), without filler degradation. This mechanism of release was found to be more effective than ion exchange. As depicted in **Figure 3F**, the release of GTM does not vary upon increasing or decreasing the concentration of cations (Na^+^) in the eluent, within the evaluated concentration range. On the contrary, CFX release is clearly affected by the concentration of anions in the elution buffer, as depicted in **Figure 3E**, further demonstrating the ion-exchange related release of the system. Compared to other inorganic anions present in dPBS, phosphate ions present the highest affinity with LDH fillers, due to their higher charge. ^[22b]^ By a 5-fold decrease in the phosphate concentration, CFX release decreased by 52% after 3 weeks. Interestingly, a 5-fold increase over the dPBS phosphate base concentration did not affect the system, suggesting the existence of a concentration threshold.

Overall, by varying the filler and antibiotic type incorporated within the scaffold, two antibiotic delivery systems were fabricated, whose sustained release can be controlled not only upon changing the filler-antibiotic concentration, but also by varying both the eluent ionic concentration or pH (in the case of MgAl-CFX), or pH (in the case of ZrP-GTM). The ionic concentration (i.e. phosphate, calcium, sodium, potassium) is a parameter that cannot be predicted during the bone regeneration process. In this regard, *in vivo* release experiments would be necessary to understand the *in situ* antibiotic release from the scaffolds. On the other hand, the local pH in a fracture during bone healing is known to be maintained in the range of pH 6 to pH 7.5 ^[35]^ and, in principle, our system is not tunable in this range. However, the property of some biocompatible and biodegradable polymers, such as the aliphatic polyesters PLA, PGA or PLGA, to dramatically decrease the local pH upon degradation (down to pH=4), ^[36]^ could be use advantageously to further customize the *in vivo* antibiotic release from filler-antibiotic composite scaffolds.

### 2.3 Thermal stability of antibiotics and scaffolds antimicrobial activity

One of the main concerns involved in the AM of 3D scaffolds via high-temperature melt extrusion of polymers containing bioactive molecules is the preservation of their bioactivity after the process’ thermal stresses. It has been previously demonstrated that the incorporation of some bioactive molecules into lamellar inorganic fillers increases their thermal stability. ^[28c, 37]^ However, CFX decomposes already at high temperatures due to its stable aromatic structure, what might have neutralized the potential shielding effect of the MgAl filler in our system, which in turn maintained CFX stability, as demonstrated by TGA (**Figure S5 A)**, as well as in previously studies. ^[37b]^ While in our TGA analysis CFX does not show any degradation bellow 255 °C, according to literature, MgAl-CFX weight loss bellow 255 °C is attributed to the loss of MgAl interlamellar water, and CFX decarboxylation ocurrs at ∼146 °C. ^[38]^ This inconsistency might be due to experimental conditions differences. Yet, HPLC confirmed molecular decomposition of CFX after the melt-blending process at around 150 °C (∼ 73% degradation), potentially attributed to the aforementioned decarboxylation of CFX at 146 °C (**Figure S5 B and C**). ^[38b]^ Further decomposition occurred during the scaffolds production via melt extrusion at 190-195 °C (∼ 45% degradation of the remaining GTM after melt blending, equivalent to ∼12.5 % degradation of the total initial GTM loading).

Similarly, GTM decomposition starts at 220 °C following some water loss, and the ZrP filler did not show to increase nor decrease its decomposition temperature, as shown in **Figure S6 A.** ^[33]^ The loss of interlamellar water of ZrP-GTM occurs in the range 20–220 °C, while the step above 220 °C is attributed to the condensation of the PO_4_^−3^ groups and the decomposition of GTM. ^[39]^ Accordingly, HPLC-MS analysis of GTM probed the stability of the molecule after each of the processing steps, with ∼ 1% and ∼ 8% degradation after melt-blending and melt extrusion AM, respectively (**Figure S6 B and D**). Interestingly, this analysis also proved the preferential intercalation of the C1a GTM isoform during the ZrP-GTM synthesis process, due to its less steric bulky form, with respect to the other isoforms (**Figure S6 C).**

Preliminary experiments demonstrated that the antimicrobial activity of the antibiotics was preserved after their intercalation within the filler, as well as in MgAl-CFX_f. Similarly, ZrP-GTM and ZrP-GTM_f presented antibiotic activity (**Table S2** and **Table S3**). Most importantly, the antimicrobial activity of both MgAl-CFX_s and ZrP-GTM_s against *S. epidermidis* and *P. aeruginosa*, was preserved on melt extrusion AM scaffold (**Figure 4**), despite the aforementioned degradation. The antimicrobial activity of CFX consists of inhibiting the action of two essential enzymes that are involved in the modulation of the chromosomal supercoiling required for bacterial DNA synthesis, transcription and cell division. ^[40]^ Despite its degradation, important active sites for CFX interaction with the target enzymes remained intact at the processing temperatures during melt-blending and posterior melt extrusion into scaffolds, thus preserving its antimicrobial activity. ^[40]^ Importantly, the antimicrobial activity of CFX contained in MgAl-CFX_s was preserved at the same levels as unprocessed CFX, observed when comparing the experimental zone of inhibition (ZOI) and the expected ZOI (i.e. ZOI given by corresponding amounts of pure unprocessed CFX) (**Table S4 and Table S5).** On the contrary, previous studies demonstrate a decrease in CFX efficiency against certain bacterial strains upon exposure to high temperatures (120 °C for sterilization). ^[19]^ However, these tests were performed under oxygen atmosphere, unlike the nitrogen conditions in which the melt extrusion AM takes place, which in combination with MgAl might have acted as a protective shield in our system. In accordance with the thermal stability results of GTM and supported by previous GTM heat stability studies ^[19]^, the efficiency of this antibiotic released from ZrP-GTM_s was also preserved at the same levels as unprocessed GTM (**Table S4 and Table S6**).

**Figure 4.**
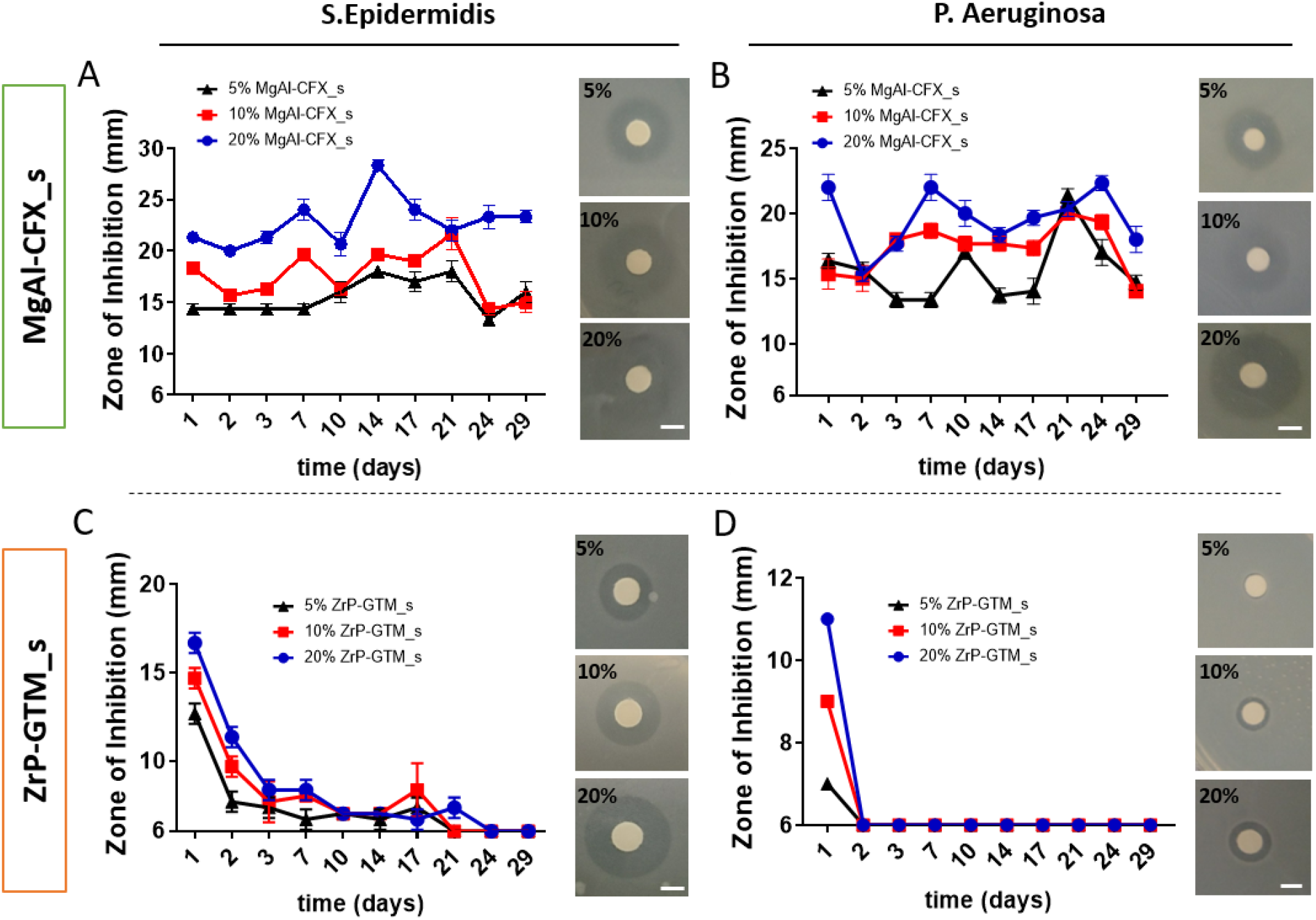
Antimicrobial activity of polymeric composite 3D AM scaffolds with different filler-antibiotic concentrations over 4 weeks. Graphs depicting zones of bacterial inhibition (ZOI) of (A, B) MgAl-CFX_s and (C, D) ZrP-GTM_s in dPBS at 37 °C against *S. epidermidis* and *P. aeruginosa*. Images of representative disk diffusion test plates depicting ZOIs around a disk impregnated with an aliquot of the scaffold release supernatant after the initial 24h of release. Scale bars 5 mm.

CFX minimal inhibitory concentrations (MIC) against both *S. epidermidis* and *P. aeruginosa* and is 1 μg/ml and 0.5 μg/ml, respectively. ^[41]^ Since the non-cumulative release of CFX from MgAl-CFX_s per timepoint was significantly above the MIC thresholds at all MgAl-CFX concentrations (**Table S4 and Figure S7**), the antimicrobial activity was preserved over the period evaluated (30 days) showing an increasing activity with increasing MgAl-CFX concentration (**Figure 4 A and B**). On the other hand, the required GTM concentration to inhibit bacterial growth is higher (MIC against *S. epidermidis*: 1 ug/ml; MIC against *P. aeruginosa* 4 ug/ml). ^[41]^ Therefore, in accordance with the amount of GTM released per timepoint (**Table S5 and Figure S7**), the antibacterial activity of ZrP-GTM_s against *S. epidermidis* lasted for 3 weeks, with a gradual decrease after a peak of activity in the first 48h (**Figure 4 C**), while only 24h against *P. aeruginosa* (**Figure 4D**). The concentration dependent activity was also observed in the case of ZrP-GTM_s s as it is shown in **Figure 4 C and D**.

While *in vivo* experiments would be necessary to fully understand the *in situ* antimicrobial function of the scaffolds, which is linked to the aforementioned *in vivo* conditions (ionic concentration, pH) for antibiotic release, this results already suggest the antimicrobial potential of the presented systems.

### 2.4 Cell viability

To further assess the cytotoxicity of the novel antibacterial systems, human mesenchymal stromal cells (hMSCs) viability was evaluated on MgAl-CFX_s and ZrP-GTM_s at the three different filler-antibiotic concentrations. Preliminary experiments demonstrated a significant reduction in cell number after 7 days of culture in 20% MgAl-CFX_s and 20% ZrP-GTM_s scaffolds (**Figure S8 A and B**). Despite their well-known biocompatibility, ^[23c, 42]^ some reports describe the dose, time and size dependent cytotoxic effects of layered lamellar compounds, such as MgAl and ZrP. ^[43]^ However, these effects were mainly seen upon particle uptake by the cells, phenomenon that it is less likely to occur during our *in vitro* experiments period as the fillers remain trapped within the copolymer matrix. Thus, it is plausible that the high amount of antibiotic release during the 4h seeding time concentrated in a small volume, negatively affected hMSCs attachment and viability. hMSCs cultured in tissue culture polystyrene showed a dose-dependence sensitivity to CFX (**Figure S9 A**), which has shown to influence MSCs proliferation upon continuous exposure. ^[44]^ Similarly, although GTM has an impact on hMSCs viability only at high concentrations (>1000 μg/ml) (**Figure S9 B**), short exposure (20 min) to high levels of GTM was found to influence osteoblasts proliferation rate up to 3 days post-exposure. ^[45]^

To prevent a decrease in cell viability, surface extraction by pre-incubating the scaffolds in cell culture media prior to seeding (for 1.5h) was considered. ^[46]^ This method was used to reduce the initial drug burst release prior to cell contact and significantly improve cell viability within our scaffolds (**Figure 5 D, E**). Accordingly, no meaningful differences in cell number were found among filler-antibiotic concentrations and comparable levels of viability were maintained for 7 days. Representative live/dead images of the scaffolds cross sections further confirmed high hMSCs viability regardless of the filler-antibiotic concentration over 7 days for both MgAl-CFX_s and ZrP-GTM_s (**Figure 5 A-C**). Interestingly, cells in the composite scaffolds showed a spread morphology comparable to the control samples (i.e. polymer-only scaffolds) after 1 day of culture, which changed to the characteristic elongated morphology of migrating cells after 7 days.

**Figure 5.**
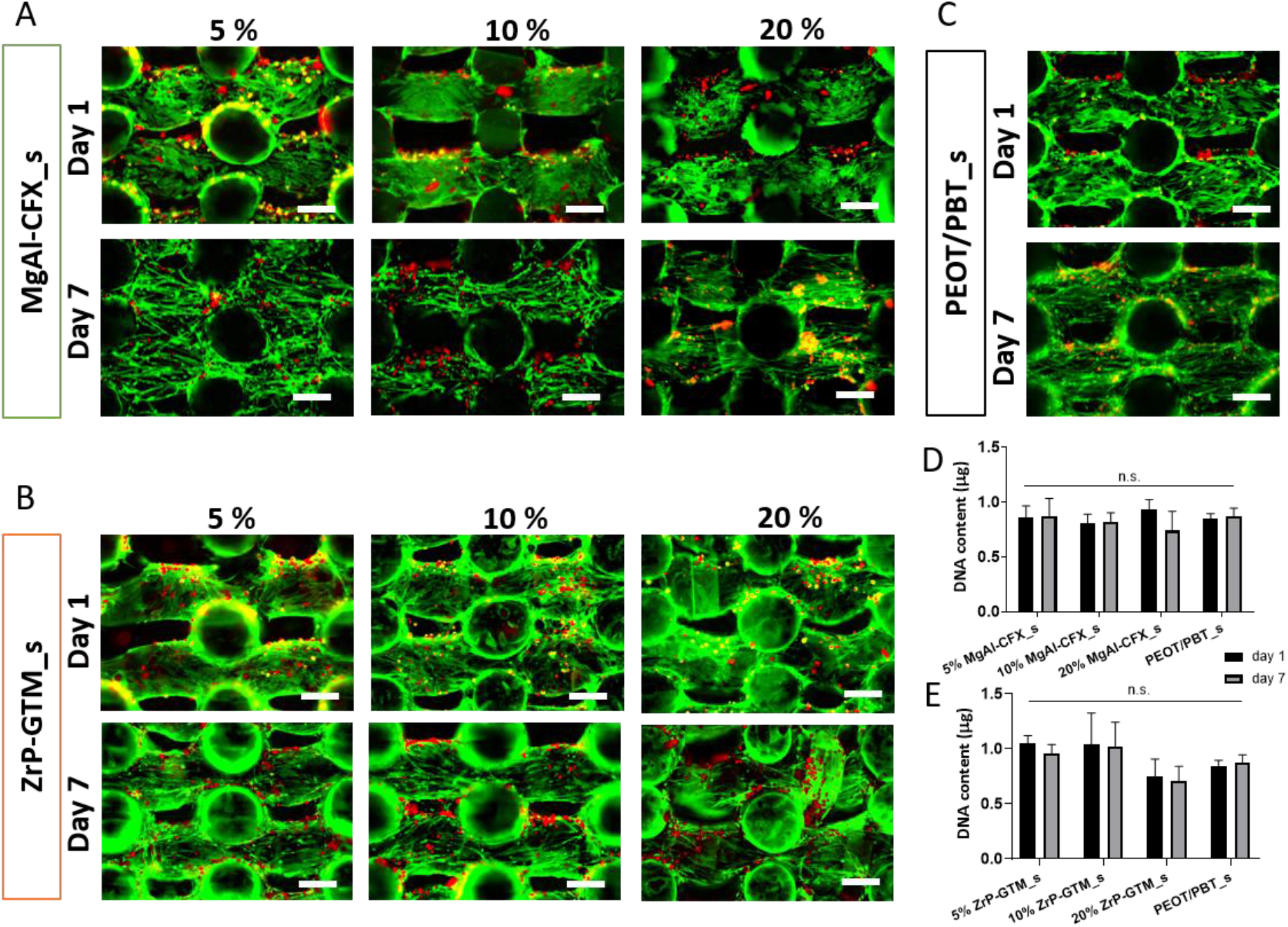
hMSCs viability is preserved after seeding MgAl-CFX_s and ZrP-CFX_s pre-incubated in media for 1.5 h. Representative fluorescent images depicting cell viability (F-actin, green; dead cells, red) 1 day and 7 days post-seeding (a) MgAl-CFX_s, (b) ZrP-GTM_s and (c) PEOT/PBT_s. Quantification of DNA content 1 day and 7 days post seeding (d) MgAl-CFX_s and (e) ZrP-GTM_s, compared to PEOT/PBT_s control. Scale bars 250 μm.

### 2.5 Osteogenic differentiation

In order to gain more insight into the potential of MgAl-CFX_s and ZrP-GTM_s to promote new bone formation, the osteogenic differentiation capacity of hMSCs seeded on the different scaffolds was analyzed. Preliminary results suggested that the osteogenic potential of hMSCs was influenced by the presence of the filler-antibiotic complexes. As depicted in **Figure S10**, after 28 days in mineralization media (MM), no mineral deposition was visualized or quantified on MgAl-CFX_s for all concentrations, nor on 20% ZrP-GTM_s, while a ∼10-fold reduction in calcium deposition was observed in 5 and 10 % ZrP-GTM_s with respect to the control samples (**Figure S10**). Yet, calcium deposits were observed in the polymer- and filler-only control scaffolds after 28 days in MM at levels reported in previous studies. ^[47]^ This indicates that the fillers were not affecting hMSCs osteogenic behavior. This is further supported by a recently published report demonstrating the significant upregulation of several osteogenic genes including alkaline phosphatase (ALP), bone sialoprotein, Runt-related transcription factor 2 (RUNX2), osteopontin (OPN), and osteocalcin in osteoblasts when adding MgAl particles to the cell culture media. ^[48]^ In order to investigate if the antibiotics were impairing or just delaying the progression of hMSCs differentiation towards the creation of a mineralized matrix, scaffolds were cultured in MM for longer periods. After a total of 35 days in MM, all ZrP-GTM_s showed calcium deposition, which were inversely proportional to the ZrP-GTM concentration (**Figure 6B**). This suggests a GTM dose-dependent effect and its role in delaying mineralization. The amount of calcium in 5, 10, and 20 % ZrP-GTM_s increased ∼3, 2 and 25-fold with respect to the previous timepoint, respectively. After a total of 49 days in MM, both 5 and 10 % MgAl-CFX_s showed matrix mineralization, which was not observed at earlier timepoints, yet at levels lower than the control conditions.

**Figure 6.**
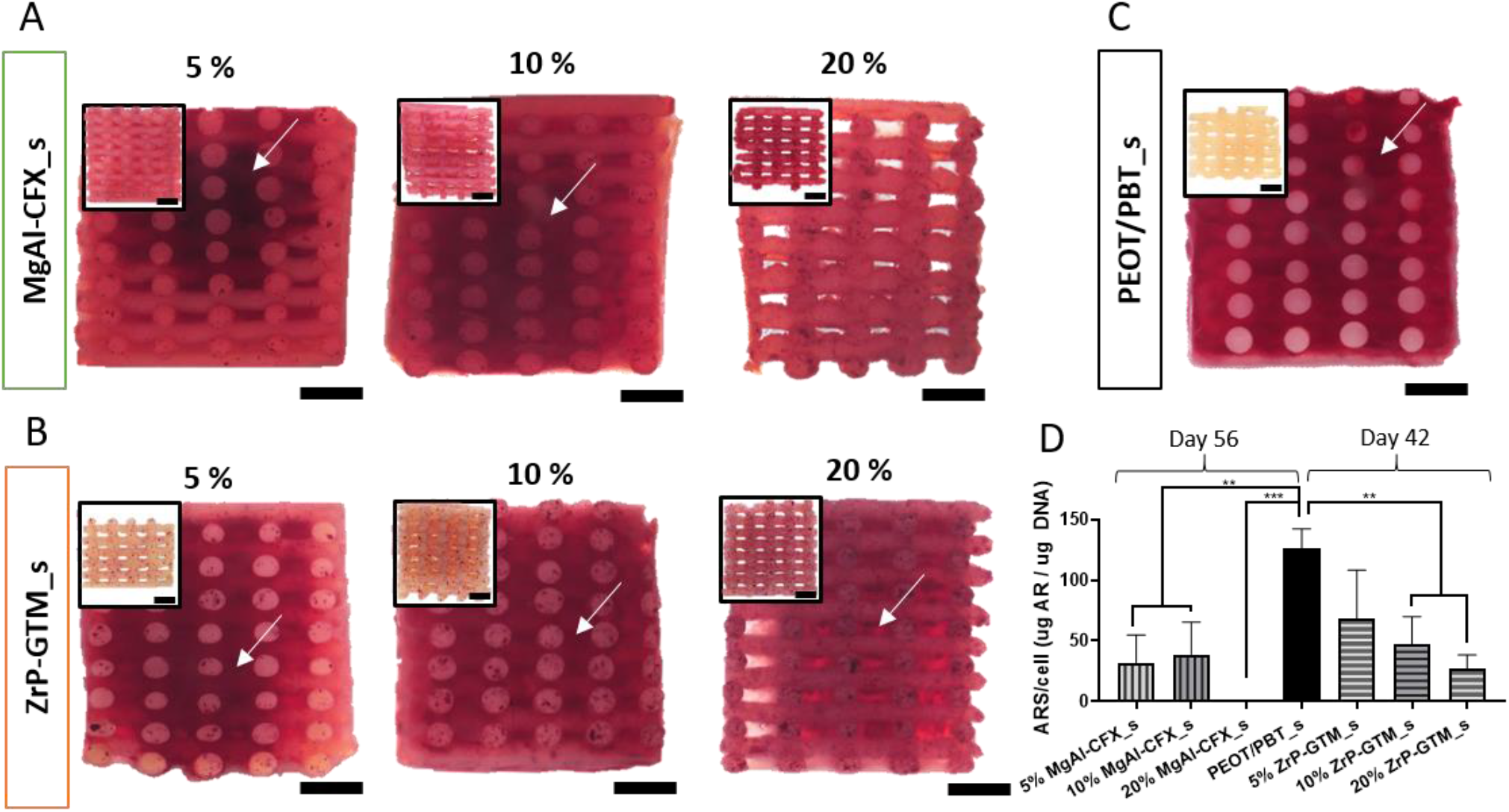
Influence of filler-antibiotic concentration on matrix mineralization. Representative stereomicroscopy images of scaffolds cross sections stained with alizarin red S: (a) MgAl-CFX_s after 56 days of culture (49 days in MM) and (b) ZrP-GTM_s and (c) PEOT/PBT_s after 42 days of culture (35 days in MM). Inserts represent the corresponding control scaffolds without cells incubated in MM and stained with alizarin red S. Scale bars 1 mm. (d) Quantification of the alizarin red S staining extracted from scaffolds normalized to cell number. Statistical significance performed using one-way ANOVA with Tukey’s multiple comparison test (**p < 0.01; ***p < 0.001).

Interestingly, 20% MgAl-CFX_s did not mineralize even after 49 days in MM, suggesting that the continuous release of higher CFX amounts had a starker effect on the hMSCs differentiation process. This result was further supported by the lower DNA and higher values of ALP activity observed on 20% MgAL-CFX_s compared to lower MgAl-CFX concentrations and the control (**Figure S11**), as the ALP activity is known to decrease during the mineralization process under optimum conditions. ^[49]^ A strong inhibition of mineralization has been reported when confluent cultures of osteoblasts were treated with CFX. The effect was apparent with 2.5-5 μg/ml CFX and progressively increased to more than a 90% decline in the calcium/protein ratio at 20-40 μg/ml CFX. ^[50]^ These values are in the range of the release of CFX from 5-10% MgAl-CFX_s and 20 % MgAl-CFX_s, respectively, therefore supporting our results. While *in vitro* experiments agree in the dose dependent interference of CFX in the bone healing process, controversial results have been reported for *in vivo* studies. Abnormalities in cartilage morphology of fracture calluses were observed after treatment in rat models with ciprofloxacin (2.4 μg/ml in serum), ^[51]^ while higher CFX release (5 μg/ml per day) from a bone defect filler in rabbits allowed bone formation within 3 months in another study. ^[52]^ This suggests that *in vivo* studies are necessary to gather a better insight over the effect of our scaffolds on bone formation. Compared to CFX, consistent *in vitro* and *in vivo* results have been previously published on the dose-dependency effect of GTM. For instance, mineralization in osteoblast and hMSCs cultures was not impaired after short exposure of up to 16 μg/ml GTM, ^[45]^ or continuous exposure of 200 μg/ml. ^[53]^ Similarly, *in vivo* GTM serum concentration of 4.5 μg/mL, ^[54]^ or a continuous release from a bone filler with a peak of 58 μg in the first 4h ^[55]^, did not show reduced healing effect in rat models. However, a burst release from a bone cement (10 mg GTM released in 3 days) resulted in significantly less mineralized tissue than the control in a sheep model. ^[56]^

Along with the study in matrix mineralization, the progression in the osteogenic differentiation of hMSCs was further evaluated by immunostaining of two relevant osteogenic makers, collagen I (COLI) and OPN. COLI constitutes nearly 90% of bone organic ECM and serves as minerals nucleation sites. ^[57]^ On the other hand, OPN is one of the most abundant non-collagenous proteins in bone. It reaches its peak during the mineralization period supporting its progression and the prevention of premature precipitation of calcium phosphate is suggested as one of its key functions. ^[49, 58]^ Interestingly, a dense layer of COLI fibrils was formed by hMSCs on top of scaffolds filaments regardless of the scaffold composition and no significant differences were observed compared to the PEOT/PBT control (**Figure 7 A and Figure S12**). This implies that COLI fibers formation was not affected by antibiotics. However, OPN expression was reduced in MgAl-CFX_s when compared to PEOT/PBT_s and ZrP-GTM_s, qualitatively observed when comparing among scaffold types the ratio of OPN-stained hMSCs over the total cell number (**Figure 7 B and Figure S12**). Intriguingly, this result lays in agreement with the delay in matrix mineralization observed on these scaffolds. Particularly, in 20 % MgAl-CFX_s, where OPN expression is more clearly downregulated, the pronounced delay in the osteogenic differentiation of hMSCs cultured on these scaffolds is confirmed once more. However, the trend of the current results indicates that longer culture periods might lead to comparable osteogenesis and matrix mineralization results among different scaffolds types.

**Figure 7.**
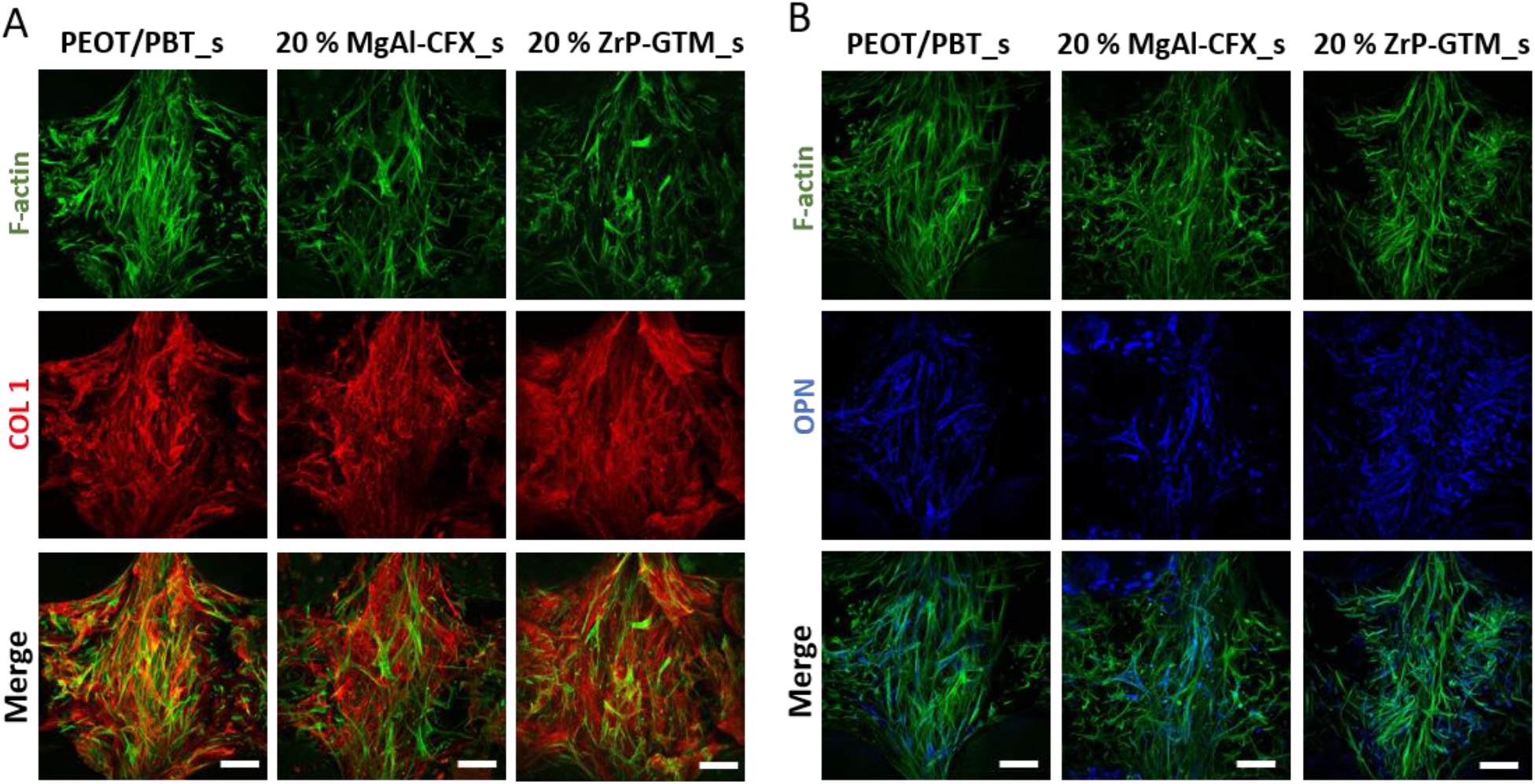
Influence of filler-antibiotic concentration on the osteogenic differentiation of hMSCs. Representative confocal microscopy images of hMSCS (F-actin, green) on top of the filaments of 20 % MgAl-CFX_s after 54 days of culture (47 days in MM) and 20% ZrP-GTM_s and PEOT/PBT_s after 42 days of culture (35 days in MM) and stained for the relevant osteogenic markers COL1 (red) and OPN (blue). Scale bars 100 μm.

Overall, the aforementioned results suggest that bone formation is supported on the filler-antibiotic based scaffolds presented in this study.

## 3. Conclusion

Local delivery of antibiotics is desired to prevent infections after bone fracture of implant-related surgery. The aim of this study was to develop antibiotic loaded 3D scaffolds fabricated via high temperature melt extrusion AM. In order to protect antibiotics from the fabrication process thermal stresses and obtain a sustained release, the antibiotics CFX and GTM were intercalated within the lamellar structure of the inorganic layered fillers MgAl and ZrP, respectively, prior to their dispersion within the polymeric matrix. Compared to no-filler systems, a more sustained release was obtained in the filler-based scaffolds. The antibiotic was released via diffusion through the polymer matrix and by a mechanism governed by ion exchange or pH changes. The functionality of the antibiotics after high temperature AM was confirmed against Gram + and Gram − bacterial strains demonstrating same antibacterial levels as unprocessed antibiotics. Importantly, the incorporation of fillers and antibiotics into the scaffolds neither affected hMSCs viability nor prevented their osteogenic differentiation, as verified by matrix mineralization and COLI and OPN expression. These newly proposed method enables the fabrication of dual functional scaffolds via direct incorporation of antibiotics in scaffolds fabricated at high temperatures, and has the potential to improve infection management while allowing bone regeneration. Moreover, this approach opens the door to future research on the use of other relevant antibiotics and biodegradable thermoplastic polymers to further optimize the system towards clinical applications.

## Supporting Information

Supporting Information is available on…..

## Acknowledgements

We are grateful to the FAST project funded under the H2020-NMP-PILOTS-2015 scheme (GA n. 685825) for financial support.

