## Supplementary Information for "3D Additive Manufactured Composite Scaffolds with Antibiotic-loaded Lamellar Fillers for Bone Infection Prevention and Tissue Regeneration"

A. Egizabal, N. Álvarez, Dr. A. Sánchez

TECNALIA, Basque Research and Technology Alliance (BRTA), 20009 Donostia-San Sebastian, Spain.

Dr. M. Bastianini, Dr. M. Sisani

Prolabin & Tefarm S.r.l., 06134 Perugia, Italy.

Dr. P. Scopece, M. Scatto

Nadir S.r.l., Mestre, 30172 Venice, Italy.

Dr. A. Bonetto, Prof. A. Marcomini

Department of Environmental Sciences, Informatics and Statistics, Ca’ Foscari University of Venice, 30172 Venice, Italy.

Prof. A. Patelli

Department of Physics and Astronomy, Padova University, 35131 Padova, Italy.

.

**Experimental section**

A list of materials described in this study and the abbreviations used to refer to them can be found in **Table S1**.

**Table S1** List of materials

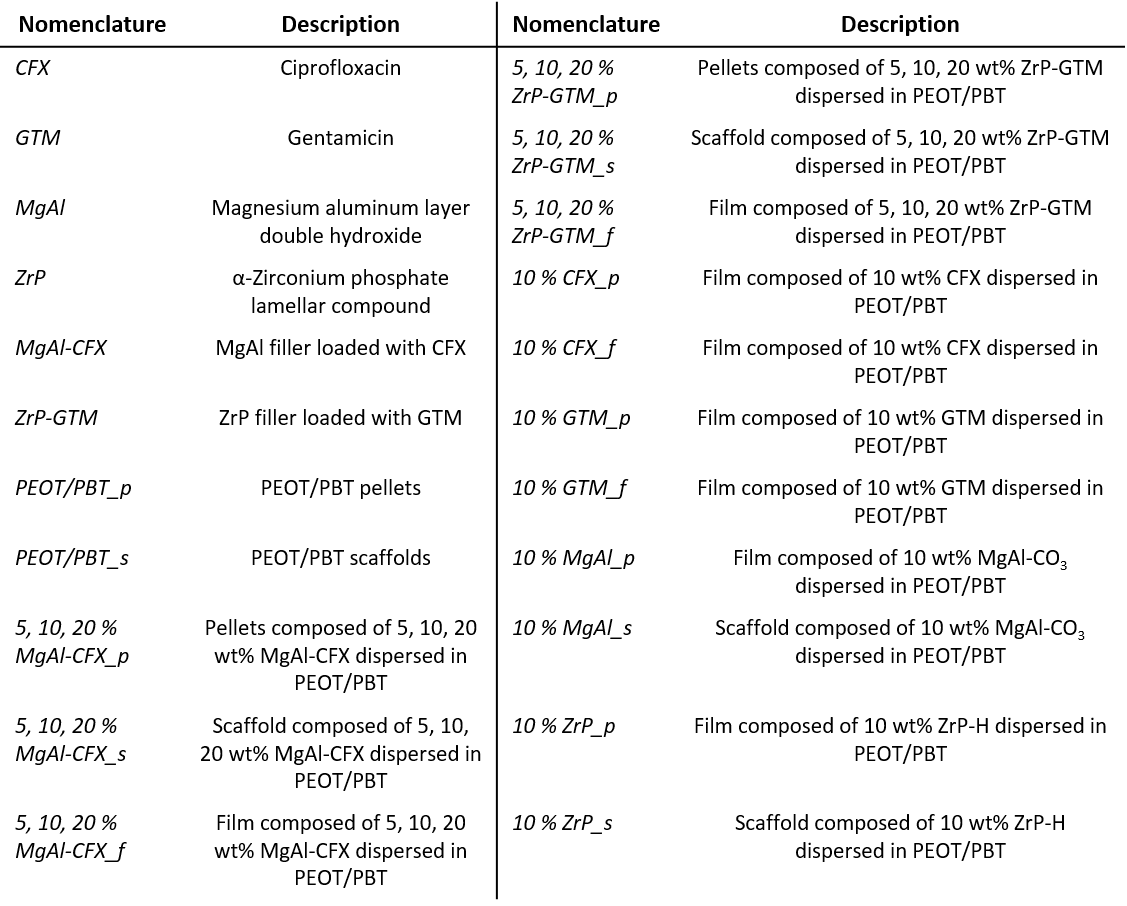

1. *MgAl synthesis and CFX intercalation*

MgAl in nitrate form (MgAl) was obtained adapting the urea method from Costantino et al. ^[1]^. Solid urea was added to 0.5 м magnesium nitrate hexahydrate (м II) - aluminum nitrate nonahydrate (м III), having molar fraction м (III)/(м (III) + м (II)) equal to 0.33, until the molar ratio urea/м (III) reached the value of 6. The mixture was heated at 100 °C for 48 h. The final product was filtered, washed with water and dried in oven at 60 °C. The intercalation compound containing ciprofloxacin (MgAl-CFX) was prepared via anionic intercalation. A carbon dioxide free water solution of NaOH 1 м (36 ml) was added to a suspension of CFX (12.2 g) in a hydroalcoholic solution (370 ml, water: EtOH 1:1) in order to obtain a sodium salt solution. Then MgAl (10 g) was added (MgAl:CFX 1:1 molar ratio). The suspension was kept under N_2_ and stirred for one day at RT. The reaction mixture was centrifuged, washed twice with a water-ethanol solution and dried in oven at 45°C.

2. *ZrP-synthesis and GTM intercalation*

ZrP synthesis and GTM intercalation was carried out as previously described. ^[2]^ Briefly, crystalline zirconium phosphate Zr(HPO_4_)2·H_2_O (ZrP) was obtained by refluxing zirconyl chloride (ZrOCl_2_·8H_2_O) in a 10 м phosphoric acid solution for 48 h. The residual solid was centrifuged, washed 3 times with water and dried in oven at 60 °C. A pre-intercalated phase of ZrP with an expanded interlayer distance, was obtained via ion exchange with propylamine, followed by HCl mixing to regenerate the acid form. The intercalation compound containing GTM (ZrP-GTM) was prepared by adding of GTM (18.9 g) to the gel. The mixture was stirred for 24 h at RT until complete exchange of the protons with GTM. The reaction mixture was centrifuged, washed twice with deionized water and dried in the oven at 40 °C.

3. *Composite production*

The production of 5, 10, 20 % MgAl-CFX_ p, 5, 10, 20 % ZrP-GTM_p, 10 % MgAl_p and 10 % ZrP_p composites was carried out in a lab scale co-rotating twin screw extruder installed in Nadir S.r.l., with a screw diameter of 11 mm and a length-to-diameter ratio (L/D) of 40. The screw profile was composed of 8 zones with three interposed kneading sections. PEOT/PBT pellets (PEOT/PBT_p, Polyvation, Netherlands) were fed in the main hopper with a volumetric feeder. Using a double inlet, MgAl-CFX or ZrP-GTM powder were fed in main hopper at 5, 10, 20 wt%. The screw rotation speed was fixed at 80 rpm while the barrel temperature was set at 140°C for the first zone and temperatures from 145 to 150°C for following zones. The nanocomposite wire was taken at the die exit, solidified in air and pelletized in a pelletizer machine. The production of 10 % CFX_p and 10 % GTM_p was carried out in co-rotating twin screw extruder by pre-mixing grinded PEOT/PBT_p and CFX or GTM powder (9:1 by weight). The screw rotation speed was fixed at 80 while the barrel temperature was set at 150°C. The nanocomposite wire was taken at the die exit, solidified in air and manually pelletized.

4. *Composite materials characterization*

TGA measurements of CFX, GTM, MgAl-CFX, ZrP-GTM, MgAl-CFX_p and ZrP-GTM_p were carried out with an STD Q600 thermal analyser (TA Instruments, USA) in air flow with a heating rate of 10 °C/min up to 800 °C. Inorganic filler content within MgAl-CFX and ZrP-GTM was calculated to be 54 wt% MgAl (46 wt% CFX) and 46 wt% ZrP (54 wt% GTM), respectively. These results in combination with the MgAl-CFX_p and ZrP-GTM_p TGA curves were used to calculate the experimental filler-antibiotic loading of the each polymer composite. Results are presented in **Table S7.**

Particle size distribution was assessed using the image analysis software tool XT Pro v3.2 (Soft Imaging System GmbH).

For HPLC analysis, a precise amount of sample was weighted. When a polymer composite was analysed, sample was dissolved in chloroform by sonication for 20 min at RT, followed by 10 min centrifugation at 4600 rpm. The supernatant was aspirated and the residual chloroform was allowed to evaporate under N_2_. The pelleted MgAl-CFX or ZrP-GTM was treated with a suitable volume of HCl (1 м) or in HCl/KCl (3 м), respectively, for antibiotic extraction. Then, the solution was filtered (pore size 0,2 µm) or centrifuged for filler separation and appropriately diluted. The amount of CFX was assessed by means of a HPLC column (Zorbax SB C18 4,6 x 250 mm, 5 µm) with a flow rate of 0.8 ml min^-1^, and the eluent consisting of water and 2% acetic acid/acetonitrile at a ratio 84/16 v/v%. In order to assess the uncertainty involved in the samples dilutions, all samples were diluted and analyzed three times. A five point calibration curve was used to quantitatively determine the CFX content.

The amount of GTM was assessed by means of a LC-MS Bruker qTOF Compact. The chromatographic separation was conducted using an Agilent Zorbax SB-C18 column (4.6 mm x 150 mm, 3.5µm) with a flow rate of 0.4 ml/min and the eluent consisting of trifluoroacetic acid (TFA) water (0.2 м) and methanol at a ratio 92/8 v/v%. The mass spectrometer was operated in full scan mode, and for the mass calibration, sodium formate clusters were used. According to literature, the four main isoform components of GTM were separated and quantified by extracting the high resolution mass chromatogram of the selected compounds with an uncertainty of 0.01 a.u.. For the quantification of the total amount of antibiotic loaded into the composites, the sum of all four species was taken into account. In order to assess the uncertainty involved in the samples dilutions, all samples were diluted and analyzed twice.

5. *Scaffolds fabrication and characterization*

Scaffolds were fabricated via melt extrusion AM. The platform (Bioscaffolder, Gesim) was equipped with a custom-made print head with two separate heating sources for the cartridge and screw. Briefly, the cartridge was filled with MgAl-CFX_p or ZrP-GTM_p, heated at 185 °C and extruded at 190-195 °C by applying a pressure of 8 bar, an auger screw rotation of 50-60 rpm and a translation speed of 15-20 mm/s. Parameters were adjusted slightly around these values for each material type and composition. 10% MgAl_p and 10% ZrP_p were also used to prepare only filler control scaffolds with the same aforementioned parameters. Similarly, PEOT/PBT_p were heated and extruded at 195 °C by applying a pressure of 4 bar, an auger screw rotation of 30 rpm and a translational speed of 15 mm/s. The scaffolds architecture consisted of a 0-90 pattern, with a 340 µm fiber diameter, 250 µm layer thickness and 850 µm strand distance (center to center), giving an expected x-y porosity of 510 µm and an expected z porosity of 90 µm. Cylindrical scaffolds of 4 mm diameter and 4 mm height were punched out from 15x15x4 mm^3^ manufactured blocks using a biopsy punch and used for further experiments.

Scaffold morphology and porosity was assessed using a stereomicroscope (Nikon SMZ25). Stereomicroscopy images were used to measure the experimental values of fiber diameter and x-y-z porosity. Presence and distribution of fillers within gold sputter-coated scaffolds’ filaments surface and cross section was examined using scanning electron microscopy (SEM, XL-30) with the backscattered electron detector (beam voltage 20 kV, spot size 5). SEM operating at 25 kV coupled with energy dispersive X-ray spectroscopy (EDS) was used to observe the chemical composition of the samples. Particle size distribution on scaffolds was assessed using the image analysis software tool XT Pro v3.2 (Soft Imaging System GmbH).

6. *Antibiotic release kinetics from films and 3D scaffolds*

To assess the antibiotic release from each of the composite 3D scaffolds, 5, 10, 20 % MgAl-CFX_s and 5, 10, 20% ZrP-GTM_s were disinfected (20 min, 70% ethanol), to mimic the disinfection performed to the samples used for *in vitro* culture, and incubated for 4 weeks in 1 ml dPBS.

2D films were prepared from 10 % CFX_p; 10 % GTM_p; 5, 10 and 20 % MgAl-CFX_p; and 5, 10, and 20 % ZrP-GTM_p. Briefly, 60 milligrams of pellets were molten at 190 °C and pressed with a coverslip against a Teflon sheet to obtain 14 mm diameter, ~300 µm thickness films. To evaluate the effect of the filler on antibiotic release, films were incubated after disinfection for 3 weeks in 1 ml Dulbecco's phosphate-buffered saline (dPBS) with both sides being exposed to the solution. To assess the effect of pH on antibiotic release, films were incubated in buffer solutions of pH 4 (potassium hydrogen phthalate based buffer), pH 7.5 (dPBS) and pH 10.5 (sodium tetraborate based buffer). To evaluate the effect of the eluent’s ionic concentration on antibiotic release, 20 % MgAl-CFX_f were incubated for 3 weeks in 1ml of dPBS (containing 10 mм Na_2_HPO_4_) or modified dPBS (with 2mм or 50 mм Na_2_HPO_4_). Similarly, 20 % ZrP-GTM_f were incubated for 3 weeks in 1ml of dPBS (containing 135 mм NaCl) or modified dPBS (with 27 mм or 685 mм NaCl).

At each timepoint, the supernatant was collected and films or scaffolds placed on 1 ml of fresh corresponding solution. CFX in the supernatants was detected by direct measurement of CFX absorbance at 275 nm using a nanodrop UV-Vis spectrophotometer (Biodrop). CFX concentrations were calculated from a CFX standard curve. GTM was detected by mixing equal volumes of supernatant, isopropanol and o-phthaldialdehyde reagent (Sigma-Aldrich) and adding 4 µl ml^-1^ 2-mecaptoethanol (Sigma Aldrich) ^[3]^. After 30 min incubation at RT, the fluorescent complex formed was detected using a spectrophotometer (CLARIOstar®, BMG Labtech) at excitation/emission= 340/455 nm. GTM concentrations were calculated from a GTM standard curve.

To investigate the drug release mechanism, 20 % MgAl-CFX_s and 20% ZrP-GTM_s release curves (% antibiotic release from the total loading vs. time) were fitted to 7 commonly used drug kinetic models: zero order, first order, Ritger-Peppas, Higuchi, Bhaskar, modified Freundlich and parabolic diffusion.

7. *Antibacterial activity*
   7.1. *Susceptibility of bacterial strains to CFX and GTM*

In order to analyze the susceptibility of bacterial strains to CFX and GTM the Agar disk-diffusion method was applied. Mueller- Hinton agar plates were spread with a standardized inoculum of *P. aeruginosa* (CECT 116) and *S. epidermidis* (CECT 231). Then, commercial filter paper discs impregnated with 20 µl of antibiotics at different concentrations (500, 250, 100, 75, 50, 25, 15, 10, 5 and 0 µg ml^-1^) were placed on the agar surface. The Petri dishes were incubated under 37 ºC during 18-24 hours. After incubation, zones of growth inhibition (ZOI) around each of the discs (including disc diameter) were measured to the nearest millimeter.

7.2. *Antibacterial activity of filler-antibiotic complexes and films*

Antibacterial activity of fillers-antibiotic complexes (MgAl-CFX and ZrP-GTM) at different concentrations (5, 10 and 20 mg/ml), and films (5, 10, 20% MgAl-CFX_f and 5, 10, 20% ZrP-GTM_f) was evaluated by shaking the samples in a concentrated bacterial suspension (106 CFU ml^-1^) in nutrient broth (1:500) for 24 h at 37 ºC. After incubation, the number of viable bacteria present in the suspension was measured by placing aliquots of the suspensions and their dilutions in sterile petri dishes with molten nutrient agar per triplicate and swirled gently. The petri dishes were incubated at 37 ºC for 24 h, after which the colonies present on the plates were counted. Values are reported as the log_10_ reduction (R), calculated as the difference in the log base 10 of the viable cell counts found on a suspension that has not been in contact with the sample and a suspension that has been in contact with the sample.

7.3. *Antibacterial activity of scaffolds*

The antibacterial activity of the antibiotics released from 5, 10, 20 % MgAl-CFX_s and 5, 10, 20% ZrP-GTM_s scaffolds incubated in dPBS solution was evaluated using the Agar disk-diffusion method, as previously described.

8. *Cell seeding and culture*

HMSCs isolated from bone marrow were purchased from Texas A&M Health Science Center, College of Medicine, Institute for Regenerative Medicine (Donor d8011L, female, age 22). Cryopreserved vials at passage 3 were plated at a density of 1000 cells cm^-2^ in tissue culture flasks and expanded until approximately 80 % confluency in complete media (CM) consisting of αMEM with Glutamax and no nucleosides (Gibco) supplemented with 10% fetal bovine serum (FBS), without penicillin-streptomycin (PenStrep) at 37 °C / 5% CO_2_.

8.1. *Metabolic activity assay to determine antibiotic toxicity*

The cytotoxicity of CFX and GTM was evaluated via PrestoBlue™ assay. Initially, trypsinized hMSCs at passage 4 were seeded at a density of 5000 cell cm^-2^ in the wells of a 48 wellplate, with CM without PenStrep, and allowed to attach for 24 hours at 37 °C / 5% CO_2_. The medium was then replaced with a fresh medium containing CFX or GTM at different concentrations, starting from 1 mg ml^-1^ or 2.5 mg ml^-1^, respectively. CM without antibiotics was used as control. After 24h, cells were washed with PBS and 250 µl of Presto Blue solution (10% PrestoBlue in CM) was added to each well and incubated for 1.5 h at 37 °C / 5% CO_2_. After incubation, the supernatants were collected and the fluorescence was measured (emission/excitation = 590/560 nm) using a spectrophotometer.

8.2. *Cell seeding on 3D scaffolds*

Scaffolds were disinfected in 70% ethanol for 20 min, washed 3 times with dPBS and incubated for 1.5 h in CM for initial burst release of the antibiotic. Scaffolds that were not pre-incubated in CM were used as controls. Before seeding, scaffolds were dried on top of a sterile filter paper and placed in the wells of a non-treated wellplate. Passage 4 hMSCs were trypsinized, centrifuged 5 min at 500 rcf and resuspended in a dextran solution (500 kDa, Farmacosmos) (10 wt/wt % dextran in CM), to achieve uniform cell seeding, at a density of 200,000 cells per 35 µl ^[4]^. The cell suspension (35 µl) was placed on top of each scaffold as a droplet, which was drawn into the scaffolds’ pores and retained inside due to the hydrophobicity of the wellplate. Seeded scaffolds were incubated for 4 hours at 37 °C / 5% CO_2_ to allow cells to attach. After this time, scaffolds were transferred to new wells containing 1.5 ml of basic media (BM) (CM supplemented with 200 μм L-Ascorbic acid 2-phosphate). BM was replaced after 24h and every two or three days from then on. To evaluate hMSCs osteogenic differentiation, after 7 days in BM, scaffolds were cultured in mineralization media (MM) consisting of BM supplemented with dexamethasone (10 nм) (Sigma-Aldrich) and β-glycerophosphate (10 mм) (Sigma-Aldrich) for another 28 days or until mineral nodules were observed (49 days in the case of MgAl-CFX_s, or 35 days in the case of ZrP-GTM_s).

8.3. *Imaging of cell viability in 3D scaffolds*

After 1h seeding and after 7 days of culture, dead cells were stained for 20 minutes prior to fixation using the Live/Dead™ fixable far-red dead cell stain kit (Thermo Fisher Scientific) at a concentration of 0.5 µl stain in 500 µl Hank's Balanced Salt Solution per scaffold. Subsequently, samples were washed, fixed with 4% paraformaldehyde for 30 min and permeabilized using 0.1% Triton X-100 for 30 min. Samples were incubated with phalloidin (Alexa Fluor 488, 1:75 in PBS) for 1h at RT as counterstain. Finally, samples were washed 3 times with PBS. Scaffolds cross sections were imaged with a fluorescent microscope (Eclipse, Ti2-e, NIKON). Background subtraction and brightness adjustments were performed on the images using the software Image J, in order to clarify their visualization. No image quantification was performed on the processed images.

8.4. *Biochemical assays*
     8.4.1. *ALP assay*

ALP activity was evaluated on MgAl-CFX_s or ZrP-GTM_s after 47 days and 35 days of culture in MM, respectively (timepoints day 56 and day 42, respectively). 3D scaffolds were collected at every timepoint and washed with PBS, stored at -80 °C and freeze-thawed 3 times, to improve lysis efficiency. Samples were incubated for 1h at RT in a cell lysis buffer composed of KH_2_PO (0.1 м), K_2_HPO_4_ (0.1 м), and Triton X-100 (0.1 v%), at pH 7.8. The chemiluminescent substrate for alkaline phosphatase CDP star® ready to use reagent (Roche) was added to the cell lysate at a 1:4 ratio. Luminescence was measured using a spectrophotometer. Remaining cell lysates were kept for DNA quantification. ALP values were reported normalized to DNA content.

8.4.2. *DNA assay*

DNA assay was performed on cells cultured on all 3D scaffolds after 1 and 7 days in BM (timepoints day 1 and day 7). Additionally it was performed on PEOT/PBT_s, 10% MgAl_s and 10% ZrP_s after 28 days in MM (timepoint day 35), on MgAl-CFX_s after 28 and 49 days in MM (timepoints day 35 and day 56), and on ZrP-GTM_s after 28 and 35 days in MM (timepoint day 35 and day 42). CyQUANT cell proliferation assay kit (Thermo Fisher Scientific) was used. Lysed samples from ALP assay or frozen samples after collection, were incubated overnight at 56 °C in Proteinase K solution (1mg/ml Proteinase K (Sigma-Aldrich) in Tris/EDTA buffer) for matrix degradation and cell lysis. Subsequently, samples were freeze-thawed three times and incubated 1h at RT with a 20X diluted lysis buffer from the kit containing RNase A (1:500) to degrade cellular RNA. Lysed samples were incubated with the fluorescent dye provided by the kit (1:1) for 15 min and fluorescence was measured (emission/excitation = 520/480 nm) with a spectrophotometer. DNA concentrations were calculated from a DNA standard curve.

8.4.3. *Alizarin red S staining and quantification*

Calcium mineralization was qualitatively determined by alizarin red S (ARS) staining on MgAl-CFX_s or ZrP-GTM_s after 28 and 49 days (timepoints day 35 and day 56, respectively), or 28 and 35 days (timepoints day 35 and day 42, respectively) of culture in MM, respectively. PEOT/PBT_s, 10% MgAl_s and 10% ZrP_s scaffolds were stained after 28 days of culture in MM (timepoint day 35). Briefly, 3D scaffolds were fixed with 4% paraformaldehyde and washed with distilled water. Subsequently, scaffolds were stained with ARS (60 mм, pH 4.1 - 4.3) for 20 min at RT, and washed with distilled water. Images os scaffolds’ cross sections were taken using a stereomicroscope (Nikon SMZ25). After imaging, stained samples (whole scaffold) were incubated for 1h at RT while shaking, followed by 10 min incubation at 85 ^o^C, with 30 v% acetic acid. Afterwards, scaffolds were removed and solutions were centrifuged at 20,000 rcf for 10 min. An appropriate volume of ammonium hydroxide (5 м) was added to the supernatants to bring the pH to 4.2. The absorbance was measured at 405 nm using a spectrophotometer. Concentration of ARS was calculated from an alizarin red standard curve and the values were normalized to DNA content.

8.5. *Immunostaining*

Cells were fixed in 4% paraformaldehyde after 35 days (PEOT/PBT_s and ZrP-GTM_s) or 47 days (MgAl-CFX_s) of culture in MM (timepoints day 42 and day 56, respectively). Triton-X 100 (0.1 v%) was added and incubated for 30 min. Samples were washed 3 times with PBS and then blocked for 1 h with blocking buffer (3 % BSA + 0.01% Triton-X 100). Afterwards, primary antibodies were added (1:200 Collagen I polyclonal antibody rabbit-derived (ab34710, Abcam), or 1:200 Osteopontin polyclonal antibody rabbit-derived (ab8448, Abcam), in washing buffer (10x diluted blocking buffer)) and incubated overnight. Samples were washed 3 times with washing buffer prior to secondary antibody incubation (1:200 Alexa Fluor 488 goat derived anti rabbit antibody (Thermo Fisher Scientific), for PEOT/PBT and MgAl-CFX scaffolds, or 1:200 Alexa Fluor 647 goat derived anti rabbit (Thermo Fisher Scientific), for ZrP-GTM based scaffolds, in washing buffer). After 1h incubation, scaffolds were washed as before and stained for F-actin (647 Alexa Fluor Phalloidin (Thermo Fisher Scientific) for PEOT/PBT_s and MgAl-CFX_s, and 568 Alexa Fluor Phallodin (Thermo Fisher Scientific) for ZrP-GTM_s (1:75 in PBS)), incubated for 1 h and washed 3 times with PBS. Confocal laser scanning microscopy was performed with a tandem confocal system (Leica TCS SP8 STED), equipped with a white light laser (WLL). Samples were excited with the dye specific wavelengths using the WLL. Emission was detected with HyD detectors (F-actin, osteopontin, collagen I). For optimal visualization in the reported images, F-actin was colored in green, collagen I in red and osteopontin in blue for all sample types.

1.1. *Statistical analysis*

Analysis of statistics was conducted with GraphPad. A one-way or two-way ANOVA was performed with a post-hoc comparison to evaluate statistical significance. Data is shown as average with error bars indicating the standard deviation.

**Supplementary Figures and Tables**

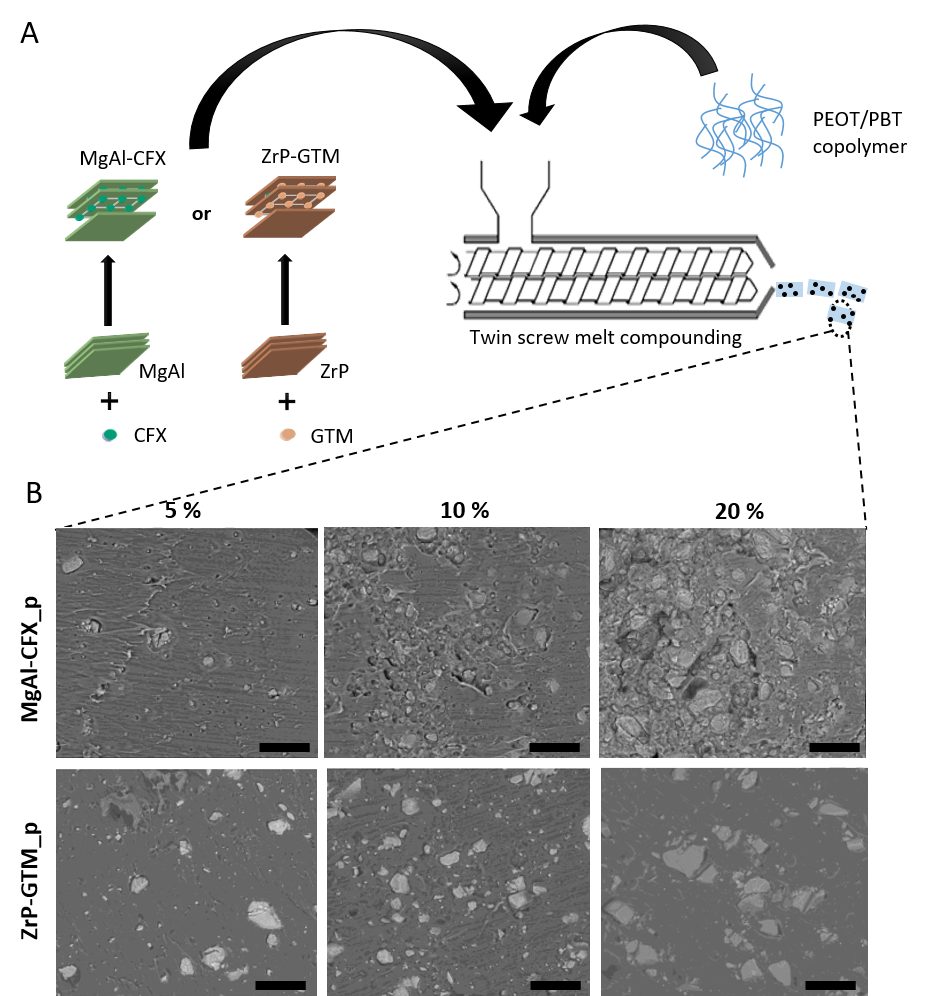

**Figure S1.** (A) Schematic representation of the melt-compounding process of the different copolymer-filler-antibiotic composites. Initially, CFX was intercalated within MgAl to obtain MgAl-CFX, while GTM was intercalated within ZrP to obtain ZrP-GTM. Each of the two filler-antibiotic complexes separately were mixed at 5, 10 and 20 wt% with PEOT/PBT using a twin screw extruder. Pellets of the 6 different materials (MgAl-CFX_p and ZrP-GTM_p) were obtained to be subsequently used for the fabrication of 3D scaffolds via melt extrusion AM. (B) SEM micrographs of the cross section of MgAl-CFX_p and ZrP-GTM_p with different filler-antibiotic concentrations. Scale bars 200 µm.

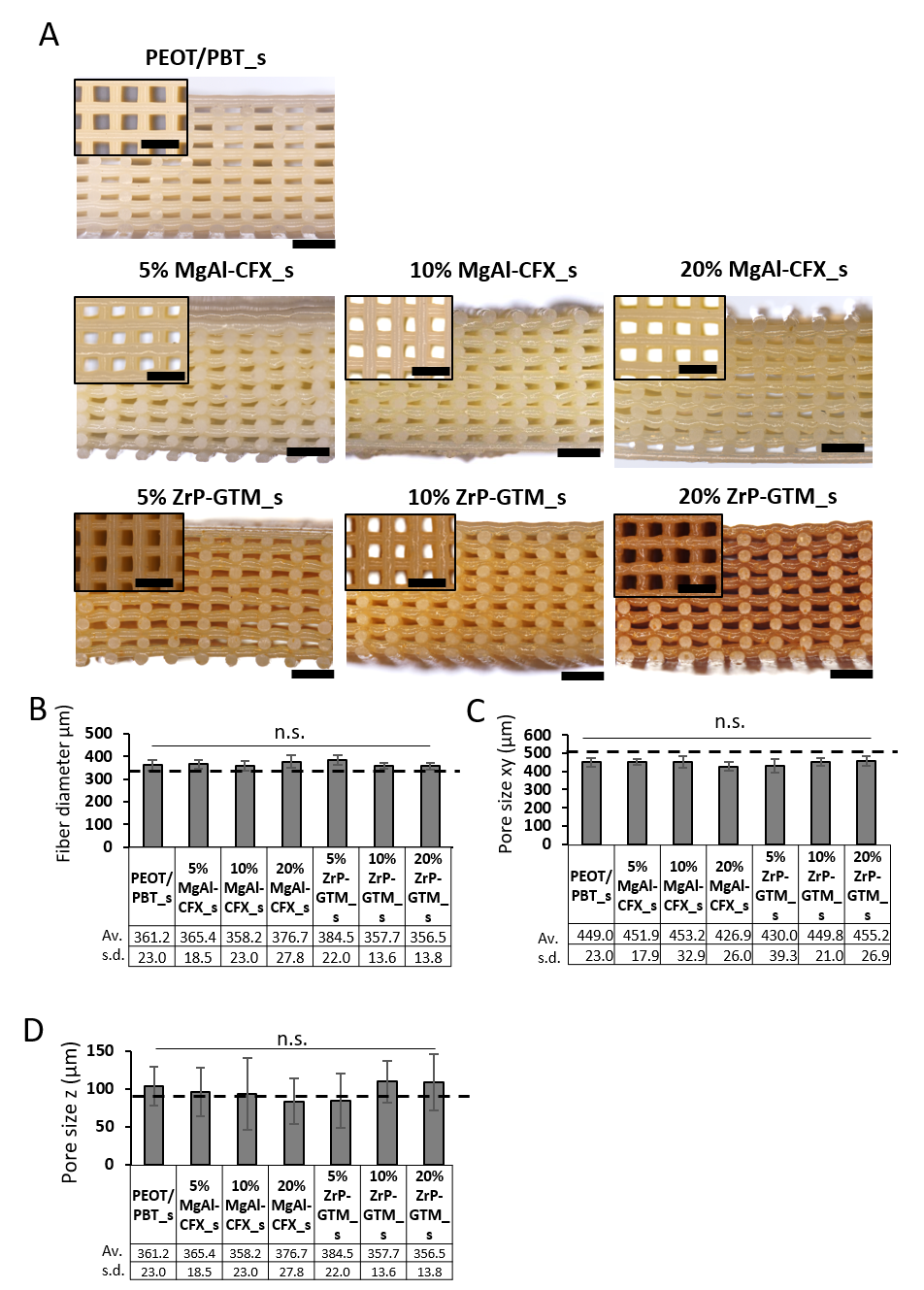

**Figure S2.** (A) Stereomicroscopy images of PEOT/PBT_s and composite scaffolds cross section and top view (inserts) depicting the scaffolds’ interconnected porosity in XYZ. Scale bars 1mm. (B) Experimental fiber diameter, (C) XY pore size and (D) Z pore size calculated for each of the scaffolds. Dash lines mark the theoretical values.

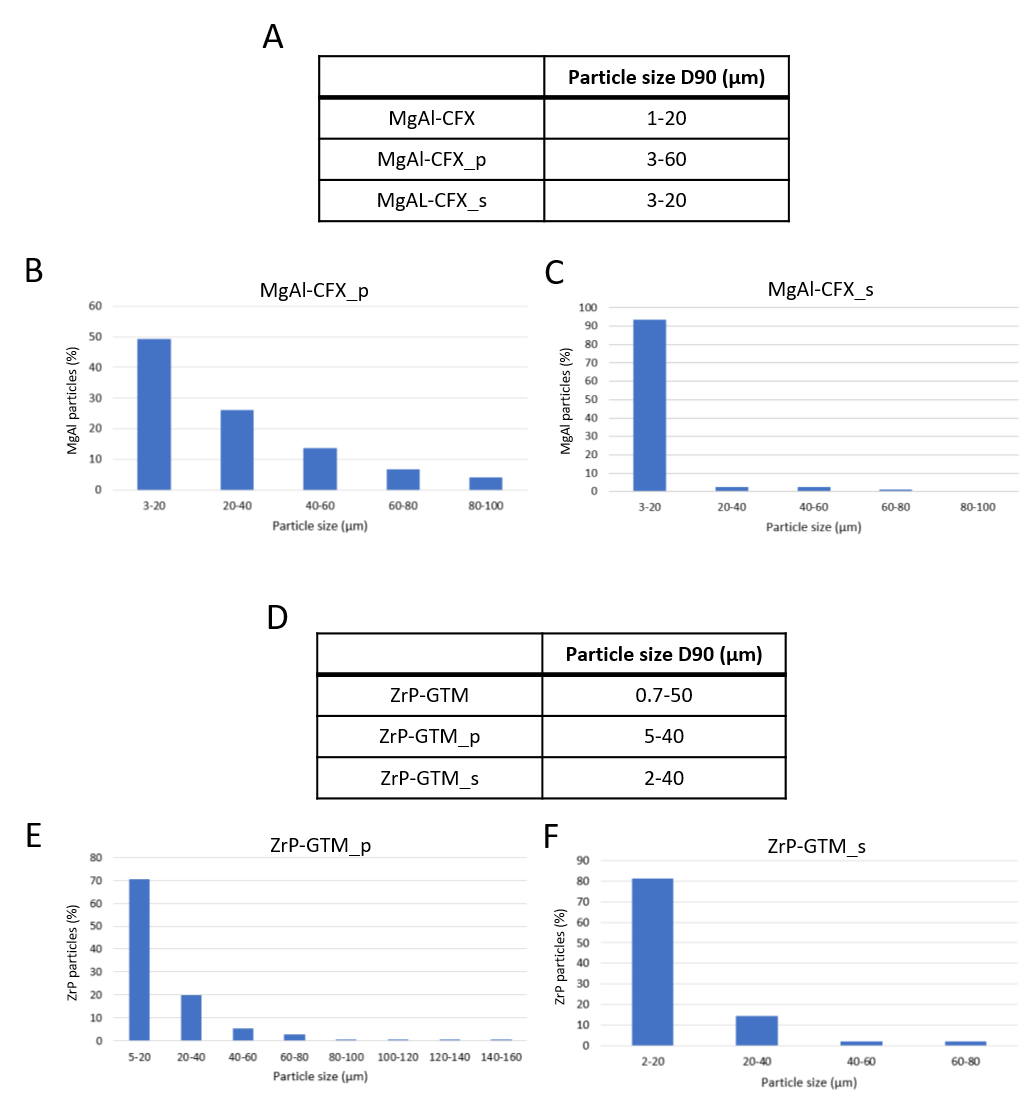

**Figure S3.** (A) MgAl-CFX particle size (D90) after synthesis, melt blending and melt extrusion AM. (B) MgAl-CFX size distribution on MgAl-CFX_p and (C) MgAl-CFX_s. (D) ZrP-GTM particle size (D90) after synthesis, melt-blending and melt extrusion AM. (E) ZrP-GTM size distribution on ZrP-GTM_p and (F) ZrP-GTM_s.

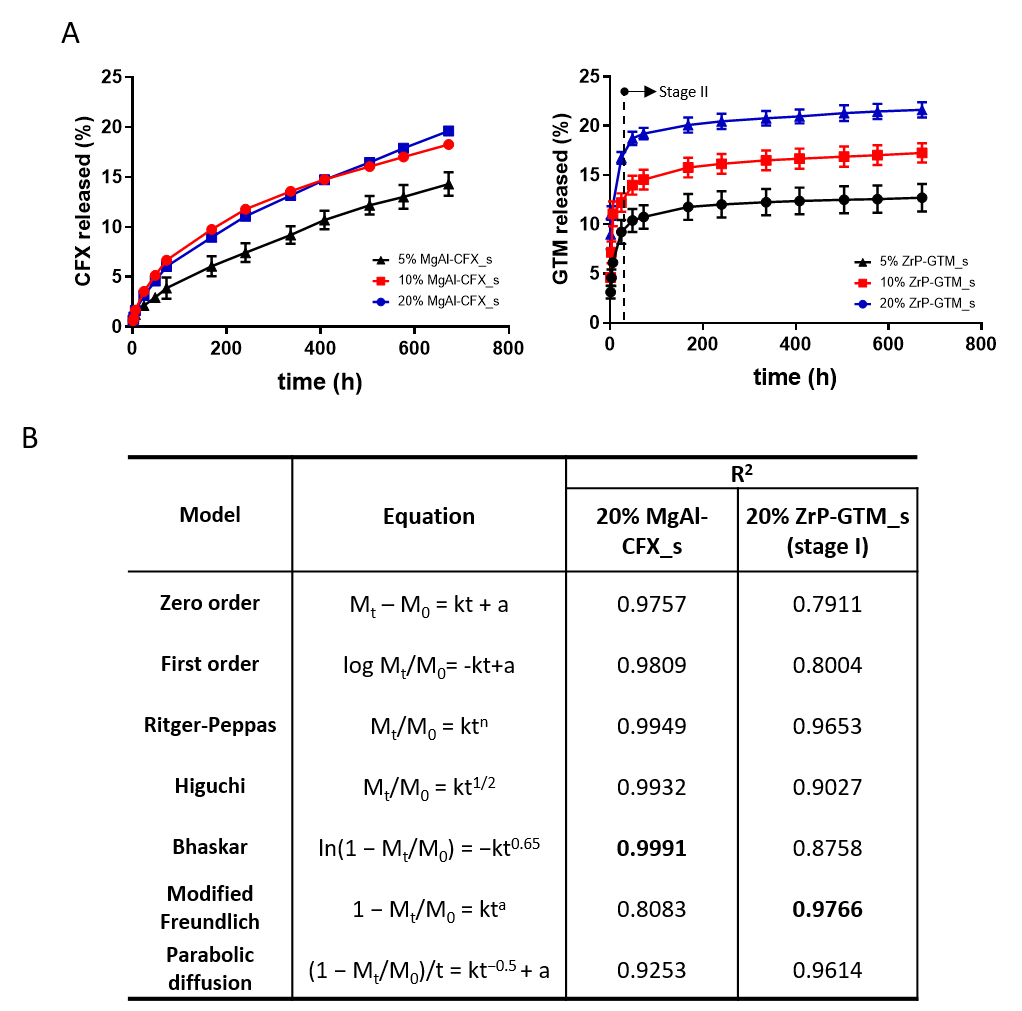

**Figure S4.** (A) Cumulative release (%) of CFX and GTM from MgAl-CFX_s and ZrP-GTM_s, respectively. (B) Kinetic models used to fit the antibiotic release profiles from 20% MgAl-CFX_s and 20% ZrP-GTM_s (stage I only) and corresponding correlation coefficients (R^2^). t refers to time, M_t_ refers to amount of drug released in time t, M_0_ refers to the total amount of drug loaded in the system, and k, a, and n are model-specific constants.

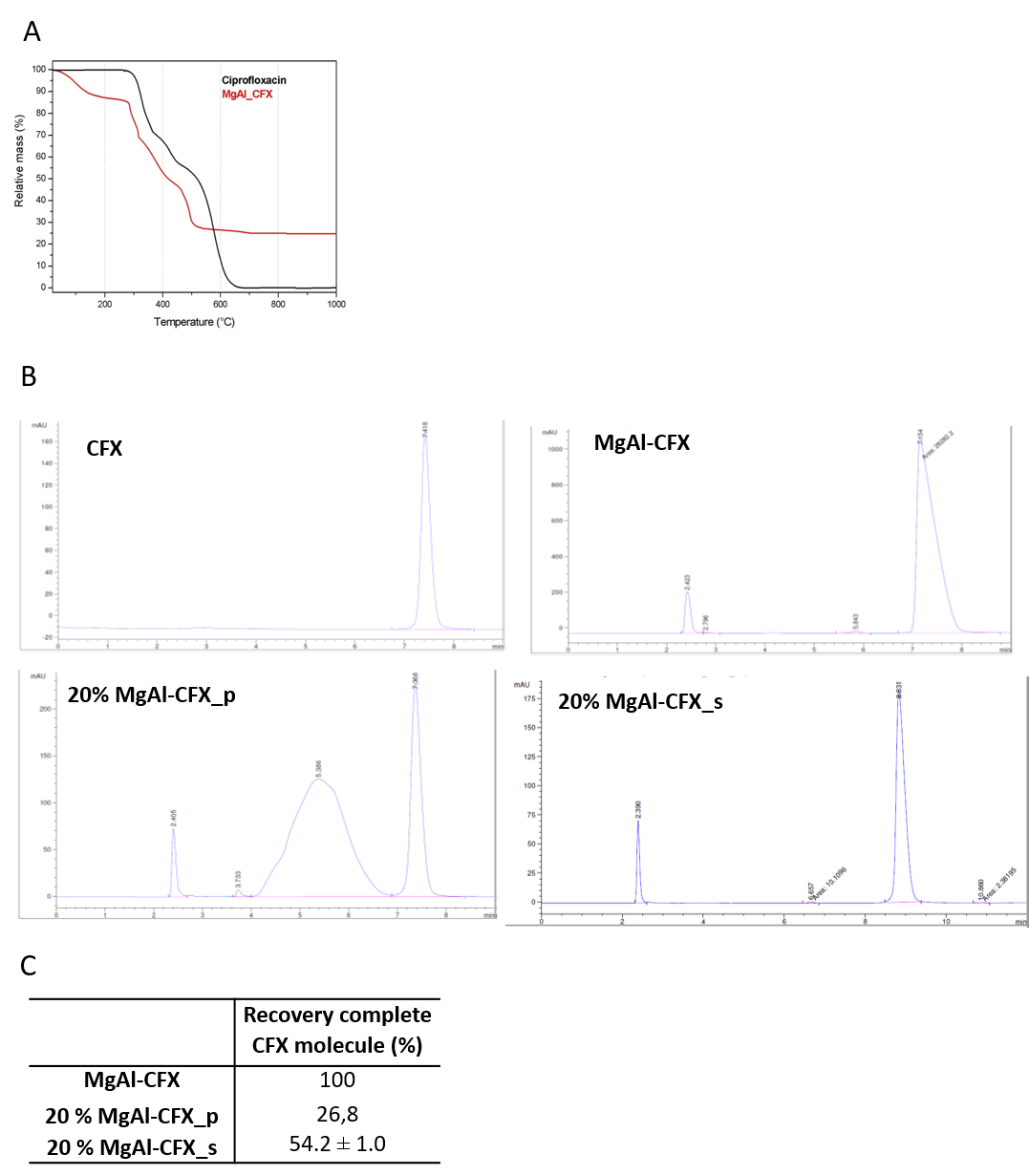

**Figure S5.** Thermal stability of CFX. (A) TGA curves of pure CFX and MgAl-CFX. (B) HPLC of pure CFX and CFX extracted from MgAl-CFX, 20% MgAL-CFX_p and 20% MgAl-CFX_s. (C) Percentage of complete CFX molecule recovery after each processing step.

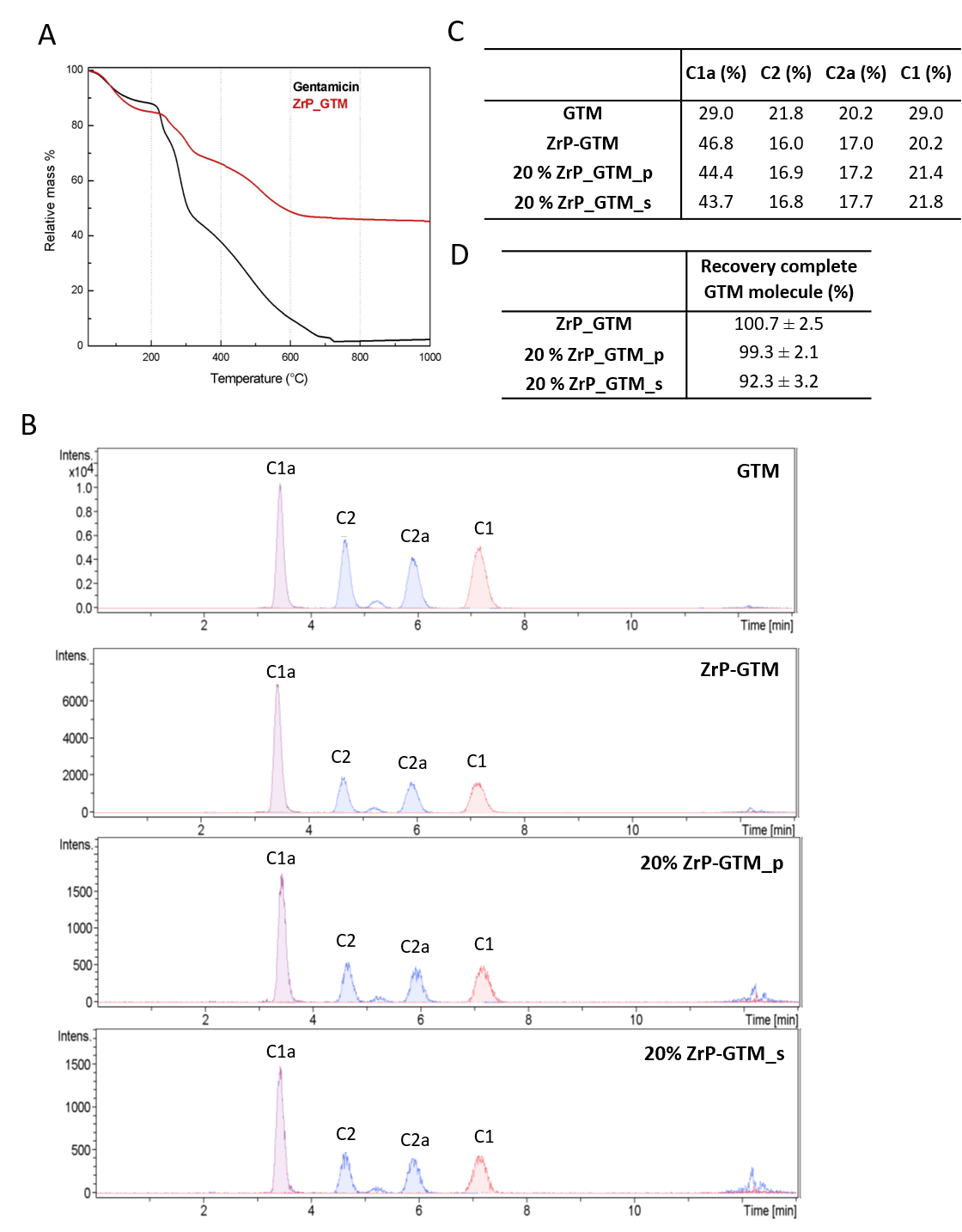

**Figure S6.** Thermal stability of GTM. (A) TGA curves of pure GTM and ZrP-GTM. (B) HPLC-MS of pure GTM and GTM extracted from ZrP-GTM, 20% ZrP-GTM_p and 20% ZrP-GTM_s. (c) Relative amount of each GTM isoform intercalated in ZrP after each processing step. (D) Percentage of complete GTM molecule recovery after each processing step.

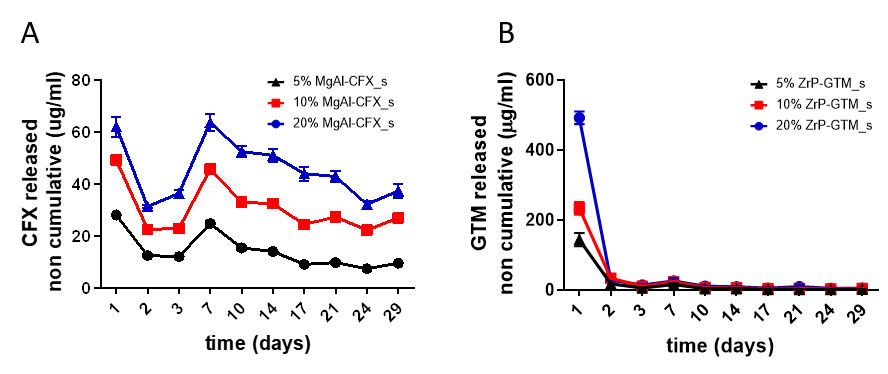

**Figure S7.** Non cumulative release of (A) CFX and (B) GTM over time from MgAl-CFX_s and ZrP-GTM_s, respectively.

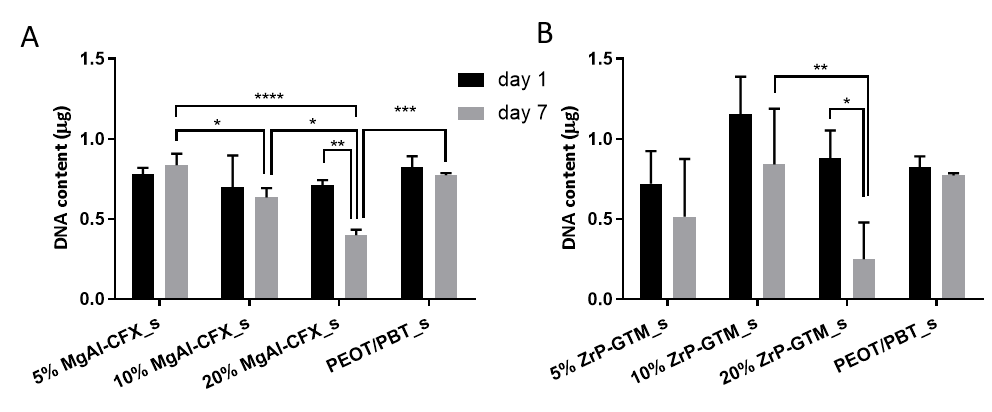

**Figure S8.** DNA content after 1 and 7 days of culture in BM in (A) MgAl-CFX_s and (B) ZrP-GTM_s, directly seeded without pre-incubation in media. Statistical significance performed using two-way ANOVA with Tukey’s multiple comparison test (* p < 0.05; ** p < 0.01; *** p<0.001; **** p < 0.0001).

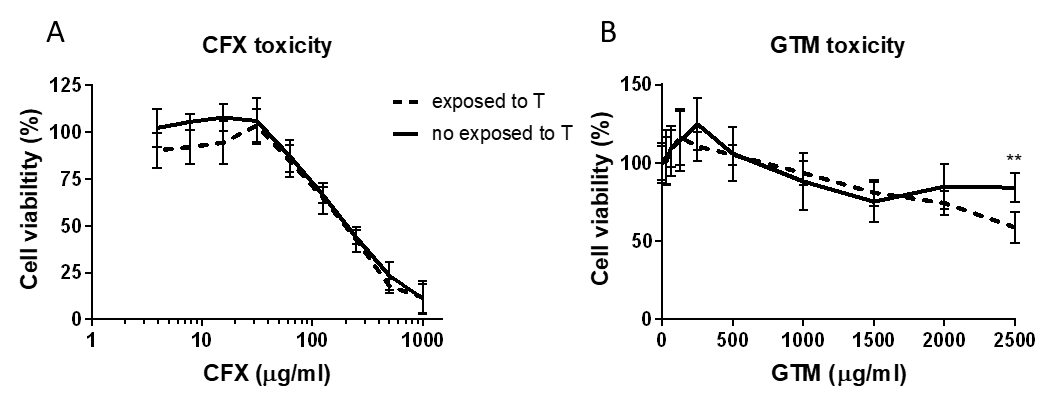

**Figure S9.** Dose dependence of toxicity of (A) CFX and (B) GTM towards hMSCs, in their pure form (not exposed to T) or previously exposed to 200 °C for 30 min under a N_2_ atmosphere (exposed to T).

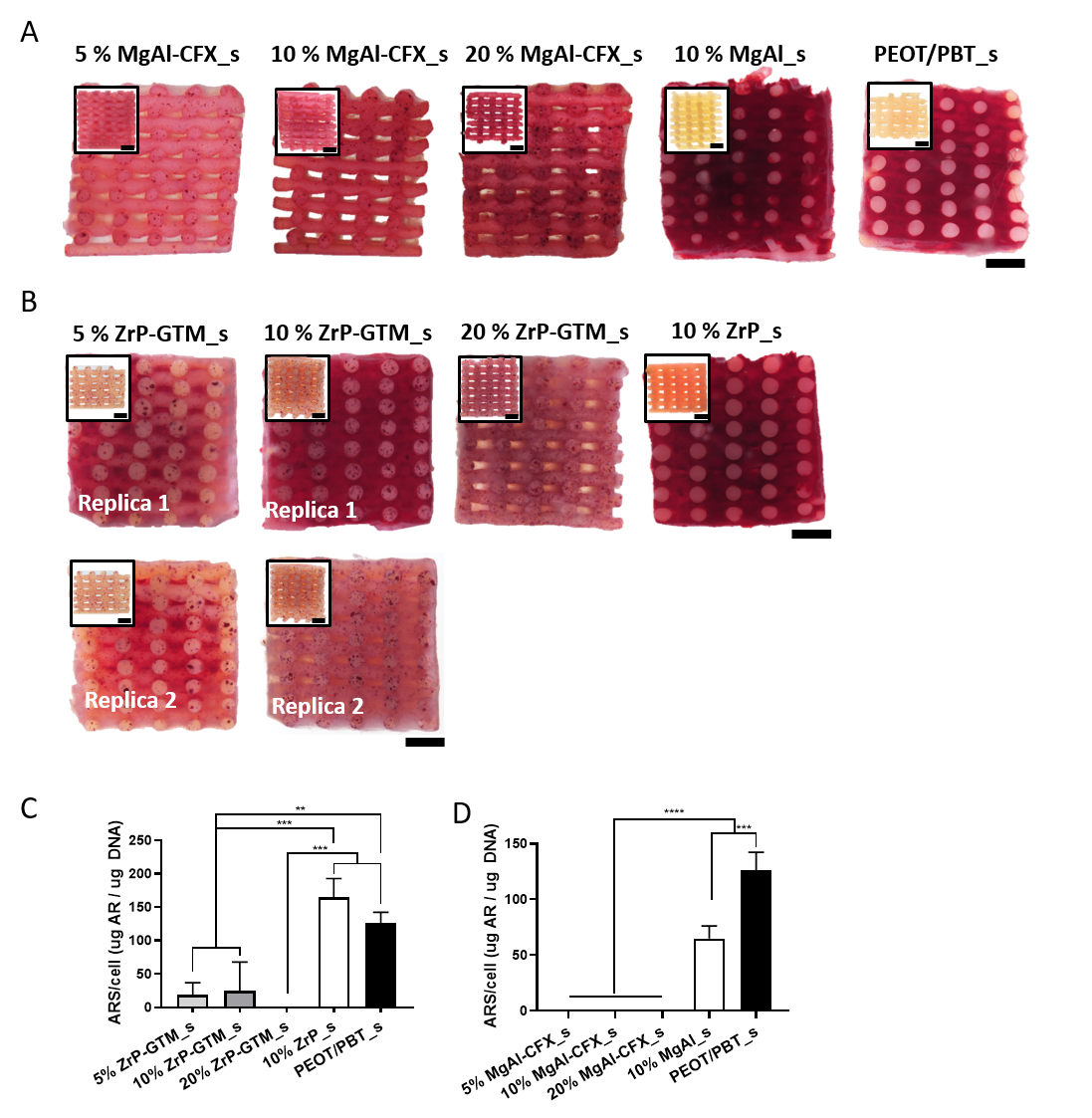

**Figure S10.** Stereomicroscopy images of (A) PEOT/PBT_s and MgAl-CFX_s and (B) ZrP-GTM_s cross sections stained with ARS after 35 days of culture (28 days in MM). Inserts represent the corresponding control scaffolds without cells incubated in MM and stained with alizarin red S. Scale bars 1 mm. (C, D) Quantification of the ARS extracted from scaffolds after 35 days of culture (28 days in MM), normalized to total cell number. Statistical significance performed using one-way ANOVA with Tukey’s multiple comparison test (**p < 0.01; ***p < 0.001; ****p < 0.0001).

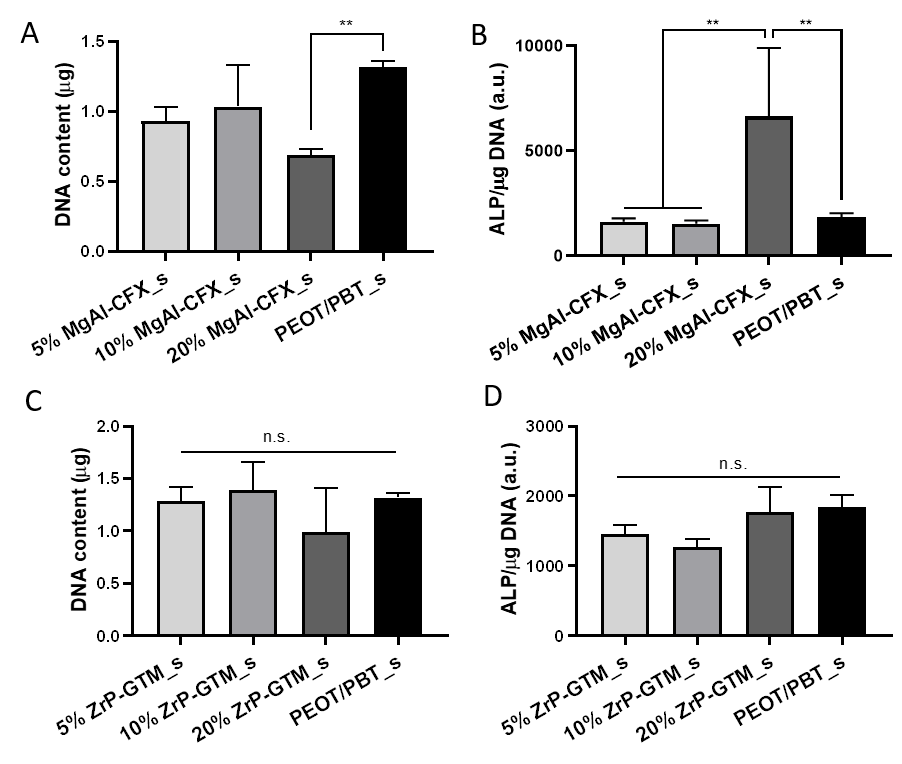

**Figure S11.** (A) DNA content and (B) ALP activity on MgAl-CFX_s after 56 days of culture (49 days in MM). (C) DNA content and (D) ALP activity on ZrP-GTM_s after 42 days of culture (35 days in MM). Statistical significance performed using one-way ANOVA with Tukey’s multiple comparison test (**p < 0.01).

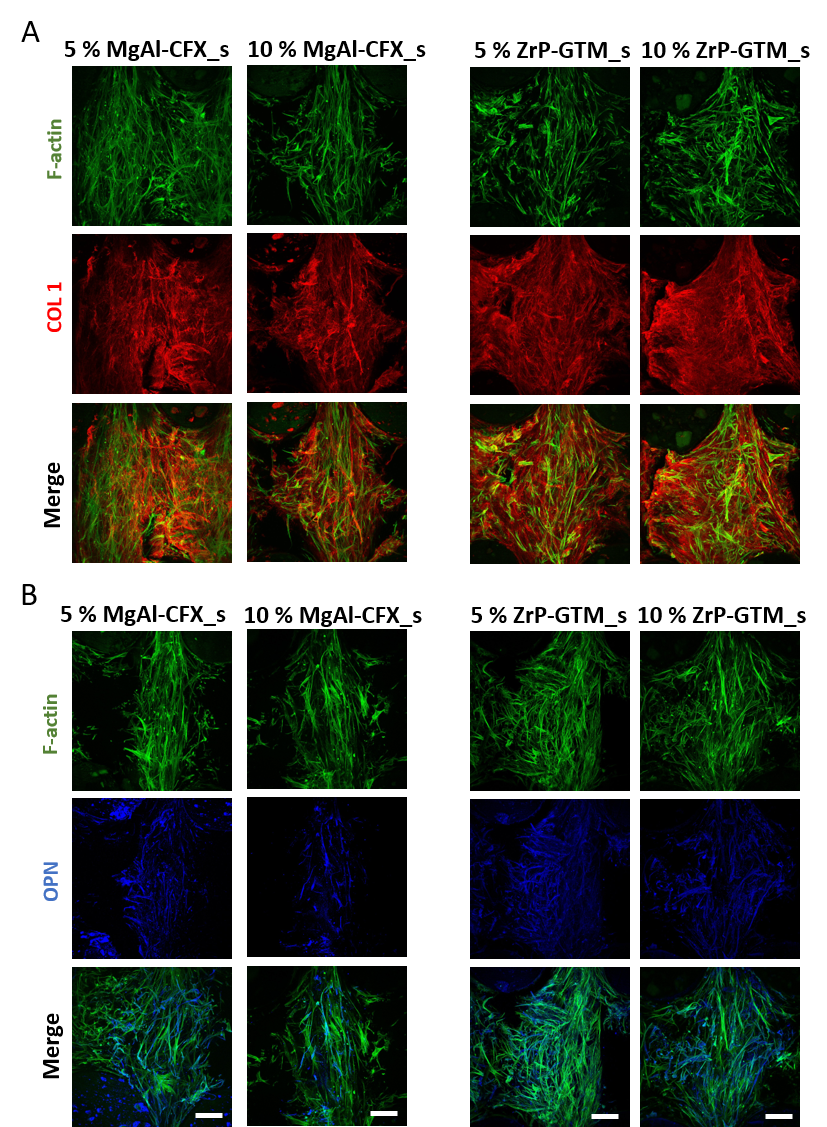

**Figure S12.** Representative confocal microscopy images of hMSCs (F-actin, green) on top of the filaments of (A) 5 and 10 % MgAl-CFX_s after 56 days of culture (49 days in MM) and (B) 5 and 10 % ZrP-GTM_s after 42 days of culture (35 days in MM) stained for relevant osteogenic markers COL1 (red) and OPN (blue). Scale bars 100 µm.

**Table S2.** Antimicrobial activity of MgAl-CFX and ZrP-GTM at different concentrations against S. epidermidis and P. aeruginosa.

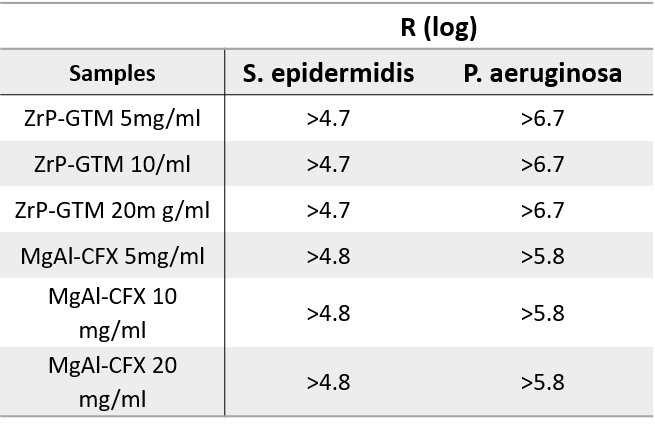

**Table S3.** Antimicrobial activity of MgAL-CFX_f and ZrP-GTM_f against S. epidermidis and P. aeruginosa.

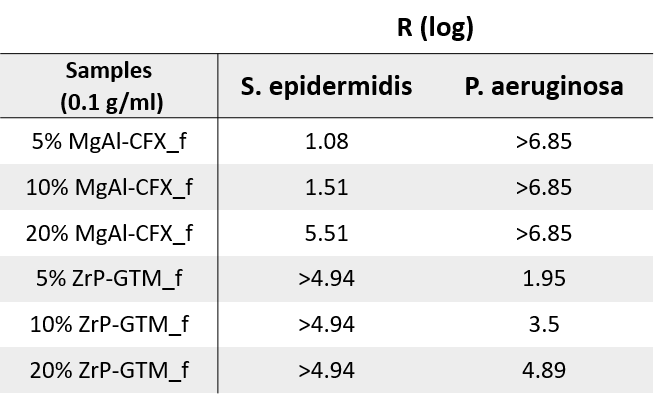

**Table S4.** Susceptibility of S. epidermidis and P. aeruginosa and strains to pure GTM and CFX evaluated by the disk diffusion agar test.

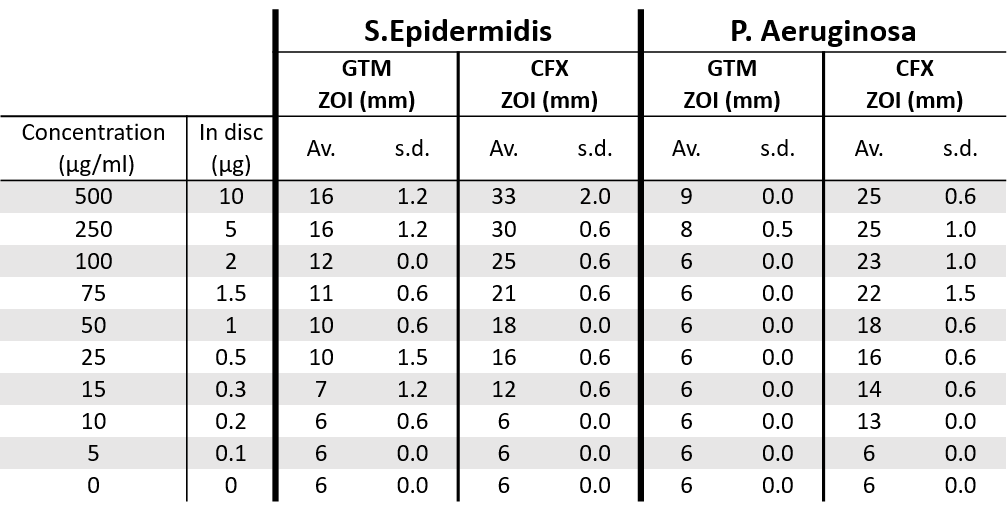

**Table S5.** Non cumulative release of CFX over time from 20 wt% MgAl-CFX_s and the corresponding experimental and expected zone of inhibitions (ZOI) against S. epidermidis and P. aeruginosa. Expected ZOI were estimated from the susceptibility test in Table S1.

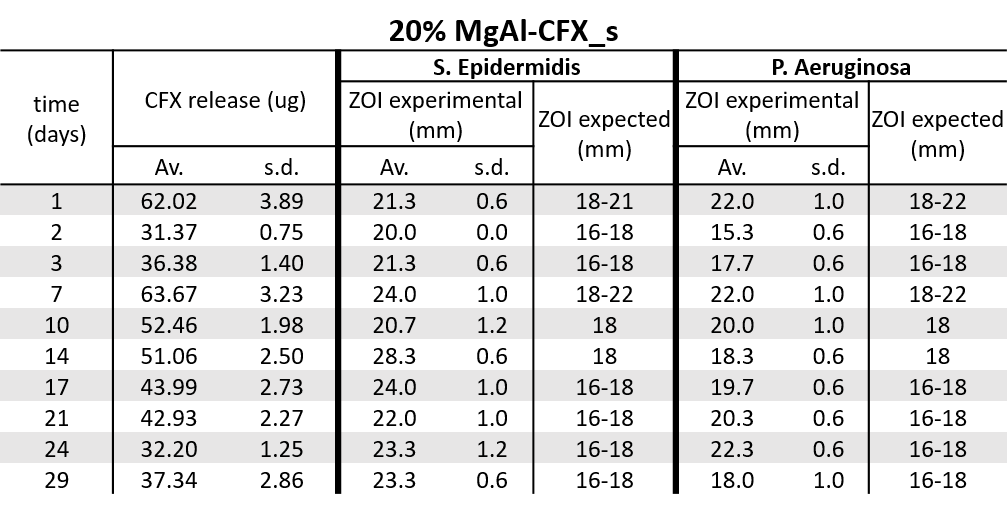

**Table S6.** Non cumulative release of GTM over time from 20 wt% ZrP-GTM_s and the corresponding experimental and expected zone of inhibitions (ZOI) against S. epidermidis and P. aeruginosa. Expected ZOI were estimated from the susceptibility test in Table S3.

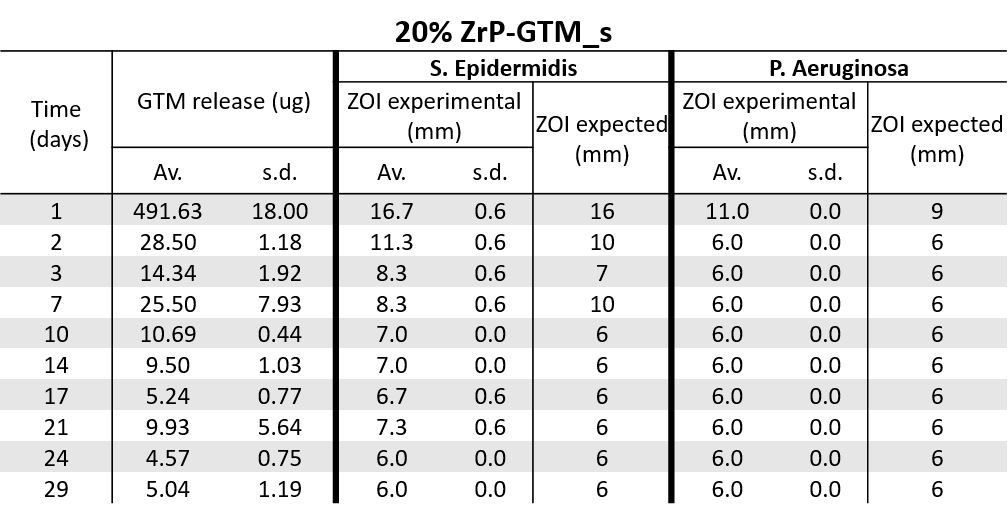

**Table S7.** Theoretical and experimental (determined by TGA) MgAl-CFX and ZrP-GTM loadings of MgAl-CFX_p and ZrP-GTM_p at different filler-antibiotic concentrations.

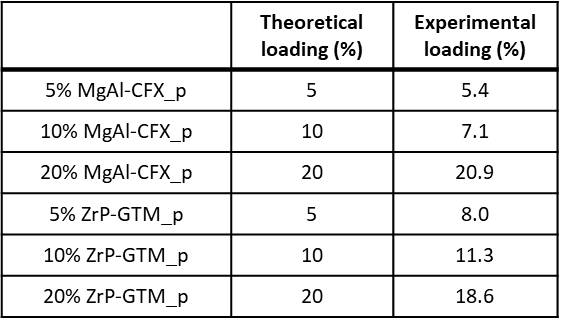
